# A High-Resolution Head and Brain Computer Model for Forward and Inverse EEG Simulation

**DOI:** 10.1101/552190

**Authors:** Alexandra Warner, Jess Tate, Brett Burton, Christopher R. Johnson

## Abstract

To conduct computational forward and inverse EEG studies of brain electrical activity, researchers must construct realistic head and brain computer models, which is both challenging and time consuming. The availability of realistic head models and corresponding imaging data is limited in terms of imaging modalities and patient diversity. In this paper, we describe a detailed head modeling pipeline and provide a high-resolution, multimodal, open-source, female head and brain model. The modeling pipeline specifically outlines image acquisition, preprocessing, registration, and segmentation; three-dimensional tetrahedral mesh generation; finite element EEG simulations; and visualization of the model and simulation results. The dataset includes both functional and structural images and EEG recordings from two high-resolution electrode configurations. The intermediate results and software components are also included in the dataset to facilitate modifications to the pipeline. This project will contribute to neuroscience research by providing a high-quality dataset that can be used for a variety of applications and a computational pipeline that may help researchers construct new head models more efficiently.

## 1 Introduction

Many simulation studies in biomedicine are based on a similar sequence of steps: image aqui-sition, creating geometric models, assigning tissue properties, performing numerical simulations, and visualizing the resulting computer model and simulation results. These steps generally describe image-based modeling, simulation, and visualization.^1–5^ Image-based modeling is useful for simulating neurological processes and can be used in applications such as electroencephalographic (EEG) forward simulation studies, EEG source imaging (ESI), and brain stimulator simulations. However, no open-source datasets or pipelines currently exist that include both functional magnetic resonance imaging (fMRI) and diffusion tensor imaging (DTI) data, which are both important in generating models of electrical propagation within the brain.

We developed a comprehensive pipeline to build a complete, high-resolution head model containing both fMRI and DTI data. The model was specifically used for EEG forward simulation studies, but can be subsequently used in ESI applications. We applied this pipeline to a healthy, female subject to develop a dataset for open-source distribution. Currently, this is the only female open-source head-modeling dataset.

In this paper, we describe every step of the pipeline, including image acquisition, preprocessing, registration, and segmentation; finite element mesh generation and simulation; and visualization. We also describe the contents of the open-source dataset, which has been released in conjunction with this paper. The open-source dataset includes raw and simulated data from the subject, intermediate results from each stage of the pipeline, and the software examples used to perform the simulations. This pipeline and dataset will be a valuable addition to the brain-modeling community because technical, resource, and expertise costs have limited the availability of such datasets.

The images, data, models, and software are available at www.sci.utah.edu/SCI_headmodel.

## 2 Methods

Our complete head model took approximately one year to complete, in part due to the many options in software and techniques, as well as the complexity of the multimodal imaging data, all of which required specific processing techniques. In this section, we describe the steps of the modeling pipeline (Figure 1). We begin with data acquisition of four image modalities followed by image preprocessing. After the images have been preprocessed, they are segmented into eight tissue layers to create a three-dimensional tetrahedral mesh. All image modalities are registered to a common coordinate space and used for forward problem simulations with processed EEG data. We describe the pipeline as applied to a female subject.

**Figure 1:**
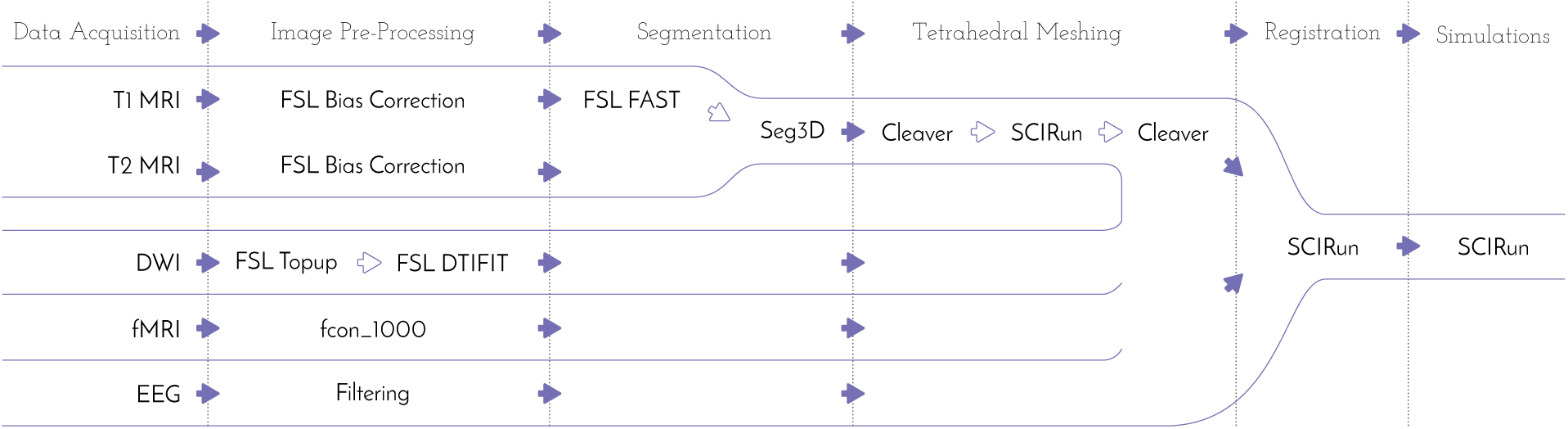
Comprehensive head/brain model pipeline. Data sources are shown on the left with software packages used for creating the head/brain model in the middle and the simulation software on the right.

### 2.1 Data Acquisition

To construct a high-resolution, personalized, anisotropic volume conductor whole-head model, *T*_1_-, *T*_2_-weighted, diffusion-weighted, and functional magnetic resonance images (MRI) were acquired on a healthy female subject, 23 years of age, on a Skyra 3T full-body scanner (Siemens Medical Solutions, Erlangen, Germany).

The *T*_1_-weighted scan was performed with a three-dimensional magnetization-prepared, rapid gradient echo (MPRAGE) sequence.^6^ The parameters used were as follows: echo time: 3.41ms; repetition time: 2500ms; flip angle: 7 °; resolution matrix size: 256×256 pixels; field of view: 256mm; 208 sagittal slices with a slice thickness of 1mm. Acquisition time was 10:42 minutes.

The *T*_2_-weighted scan was performed with a SPACE (sampling perfection with application)-optimized contrast using different flip angle evolutions – sequence.^7^ The parameters used were as follows: echo time: 406ms; repetition time: 3200ms; resolution matrix size: 256×256 pixels; field of view: 256mm; 208 sagittal slices with a slice thickness of 1mm. Acquisition time was 5:34 minutes.

The diffusion-weighted images (DWI) were acquired with multiband, two-dimensional, echo-planar imaging (EPI).^8^ Both phase-encoding directions were performed (anterior to posterior (AP) and posterior to anterior (PA)) with 64 diffusion directions each. Further sequence parameters for each scan were as follows: echo time: 76.8ms; repetition time: 4070ms; flip angle: 90 °; resolution matrix size: 104×104 pixels; field of view: 208mm; 60 slices with 2.5mm slice thickness. Acquisition time was 5:05 minutes each.

The functional MRI (fMRI) scans were acquired with a blood-oxygenation-level dependent contrast (BOLD) sequence. The following parameters were used: echo time: 76.8ms; repetition time: 780ms; flip angle: 55 °; resolution matrix size: 104×104 pixels; field of view: 210mm; 72 slices with 2mm slice thickness. Acquisition time was 10:32 minutes.

Continuous electroencephalograms (EEGs) were recorded using a 128-channel and 256-channel HydroCel Geodesic Sensor Net that was connected to a NetAmps 400 amplifier and referenced online to a single vertex electrode. Channel impedances were kept at or below 50 kOhms, and signals were sampled at 250Hz. The EEGs were recorded while the subject sat quietly in a chair, alternating 2 minute epochs of eyes open and eyes closed, for a total of 12 minutes.

All acquisition reports are included with the dataset.

### 2.2 Preprocessing of Images

#### 2.2.1 MRI Correction

A bias field signal is a low-frequency, smooth signal that corrupts MRI images due to inhomogeneities in the magnetic fields of the MRI machine by blurring images, thereby reducing the high frequencies of the images, such as edges and contours. The signal changes the intensity values of image pixels so that the same tissue has a different distribution of grayscale intensities across the image.^9^ We applied an estimated bias field correction on the *T*_1_ and *T*_2_ MRIs using FMRIB Software Library (FSL) FAST (Figure 2),^10^ which will be further described in Section 2.3.

**Figure 2:**
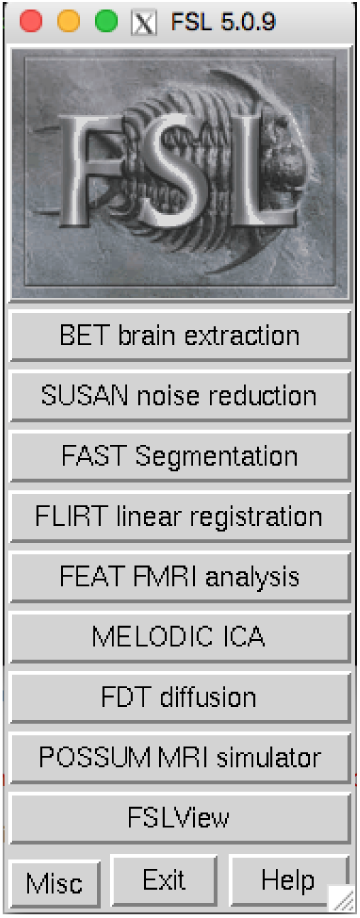
FMRIB software library user interface.

#### 2.2.2 DWI Distortion Correction

DWIs performed with EPI sequences are prone to distortions from rapid switching of diffusion-weighting gradients, movement from the scanning table, and movement from the subject. The diffusion data were collected with reversed phase-encoded blips (AP and PA), resulting in pairs of images with distortions in opposite directions. From these pairs, we estimated the susceptibility-induced off-resonance field using a method^11^ similar to what is currently implemented in FSL.^12^ We then combined the two images into a single corrected image using FSL’s topup and eddy command line tools. Details on this process are included in the Appendix in Section A.1.

#### 2.2.3 Diffusion Tensor Images

After we preprocessed the DWI images, we calculated diffusion tensor images (DTI) using FSL’s DTIFIT toolbox^13^ (Figure 3) and SCIRun v4.7,^14^ a problem-solving environment for modeling, simulation, and visualization of scientific problems. We used the output from DTIFIT, the eigen-values and eigenvectors of the DTI data, as input for a SCIRun network (Figure 36) to build and visualize the conductivity tensor field. Details on using DTIFIT are included in the Appendix in Section A.2.

**Figure 3:**
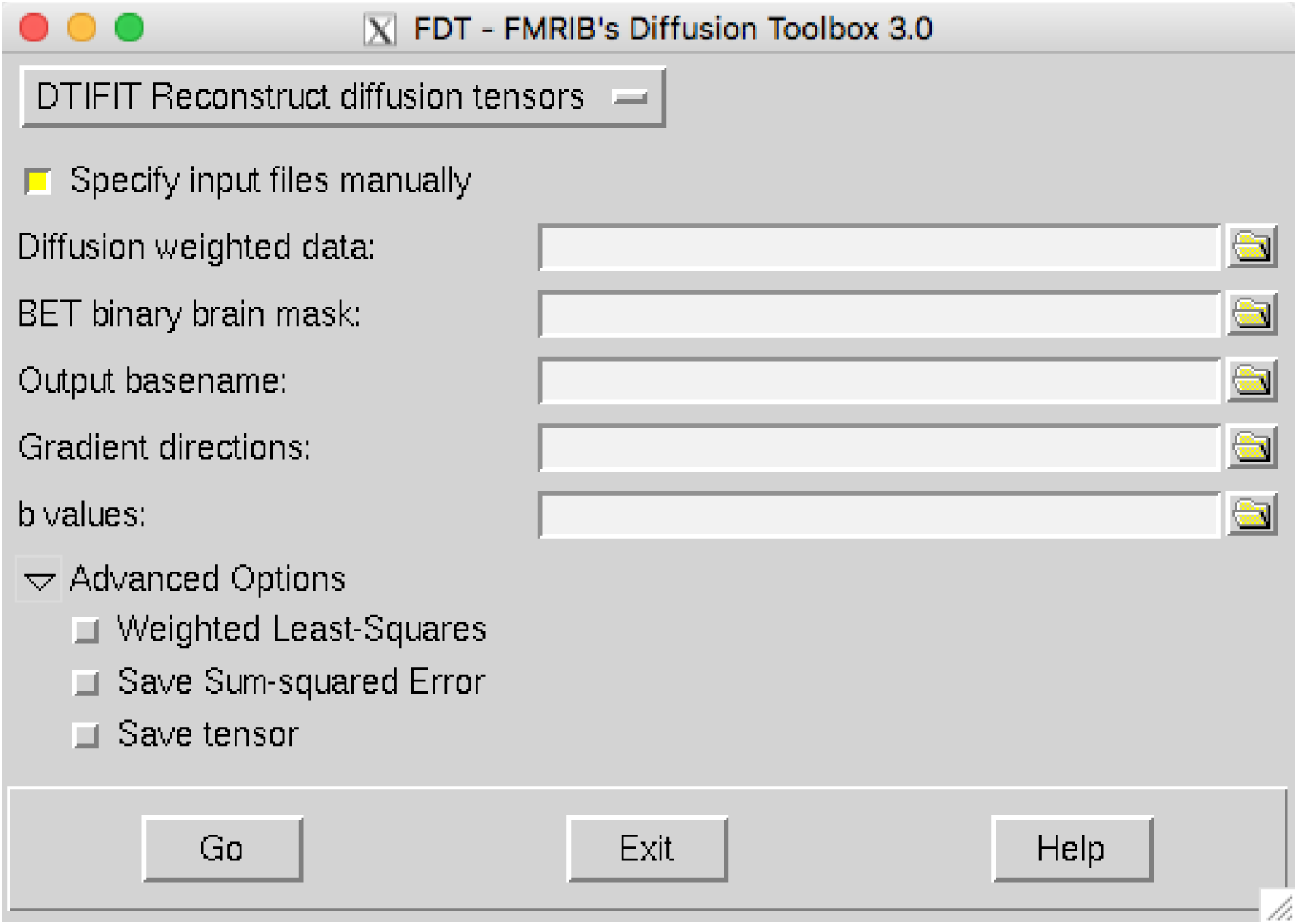
FSL DTIFIT user interface.

#### 2.2.4 fMRI

We preprocessed the fMRI data using the 1000 Functional Connectomes (fcon) Project pipeline scripts,^15^ which performed anatomical preprocessing, functional preprocessing, registration to the *T*_1_ MRI, segmentation, and nuisance signal regress. The outline pipeline used on this fMRI dataset, specific to the University of Utah, can be found at https://bitbucket.org/UtahBrainNetworksbase_prep, which includes instructions for installation, compilation, and usage. We then converted the preprocessed fMRI data from a four-dimensional to a two-dimensional dataset to be visualized in SCIRun (Figure 39). Details on this data conversion are included in the Appendix in Section A.3.

#### 2.2.5 EEG

We applied a 60Hz notch filter and its harmonics^16^ to the EEG data to create an EEG data matrix. The rows of the matrix corresponded to the channels (electrodes) of the EEG net and the columns corresponded to the time step. We removed the last two rows of the EEG data matrix, because these were controls for the experiment, and several columns at the beginning and end of the matrix, because they corresponded to taking the EEG net on and off the subject’s head. Details on building the EEG data matrix are included in the Appendix in Section A.4.

#### 2.2.6 Registration

Since the subject did not move in between the *T*_1_ and *T*_2_ MRI, registration was not necessary before segmentation and mesh generation. We generated the tetrahedral mesh from the segmentation and registered the cortical surface mesh to the DTI coordinate space with a manual, rigid registration (Figure 4 – *(left)*). We included the registration transformation matrix to the DTI coordinate space in the dataset.

**Figure 4:**
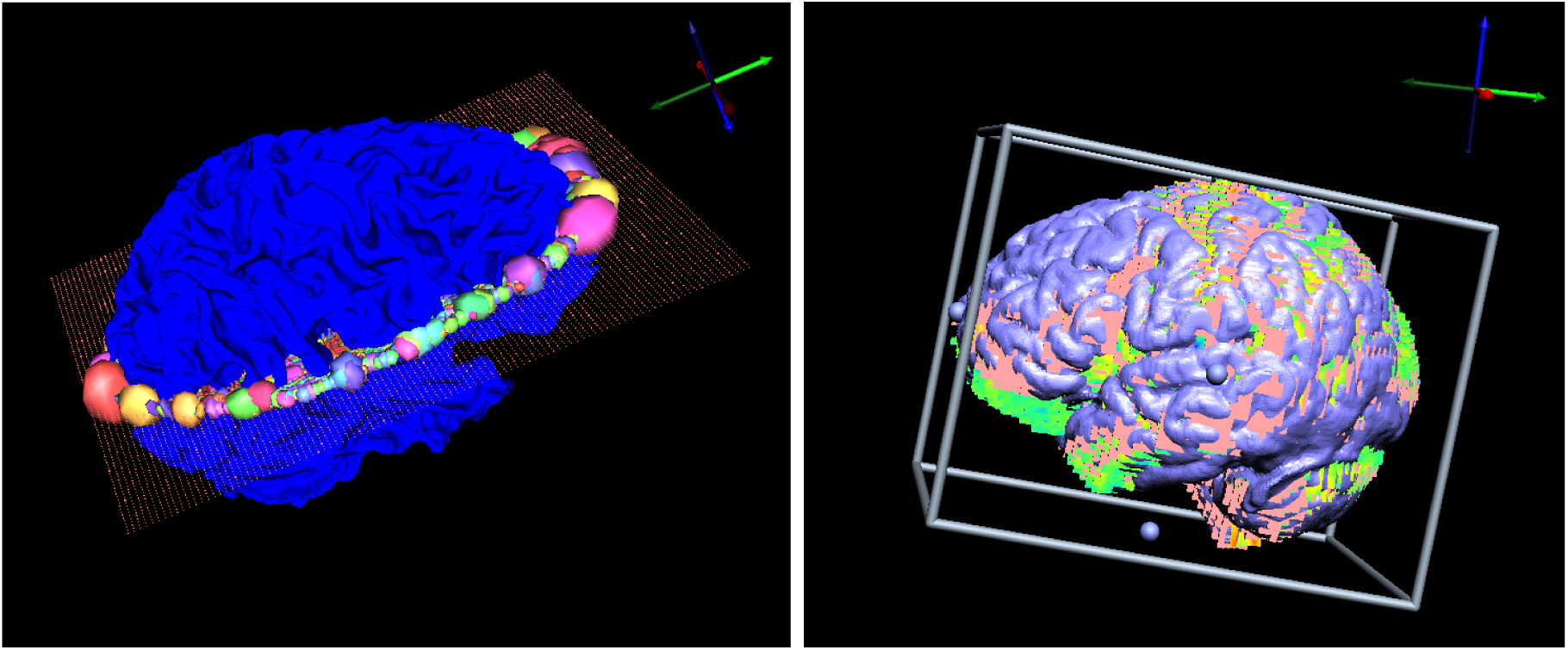
SCIRun manual registrations: cortical surface mesh to DTI registration *(left)*, fMRI to cortical surface mesh registration *(right)*.

Registration of the fMRI dataset required spatial preparation to ensure a more accurate rigid registration. For each fMRI time step, we mapped the corresponding vector of fMRI data onto a lattice volume, rotated the volume 180 degrees about the *z*-axis, smoothed it with a median filter, applied a threshold to it to remove noise, and clipped the brainstem. After we applied the rigid registration to the cortical surface mesh, we continued the registration manually to ensure the most accurate registration (Figure 4 – *(right)*).

The EEG geometry data from the ESI system consisted of dipole source locations (4800) and electrode locations (128 or 256). To register the electrode positions to the head mesh, we rotated them 180 degrees about the *z*-axis, applied a rigid registration, and projected the points onto the head surface mesh (Figure 5). We then applied a rigid registration to register the dipoles to the cortical surface mesh.

**Figure 5:**
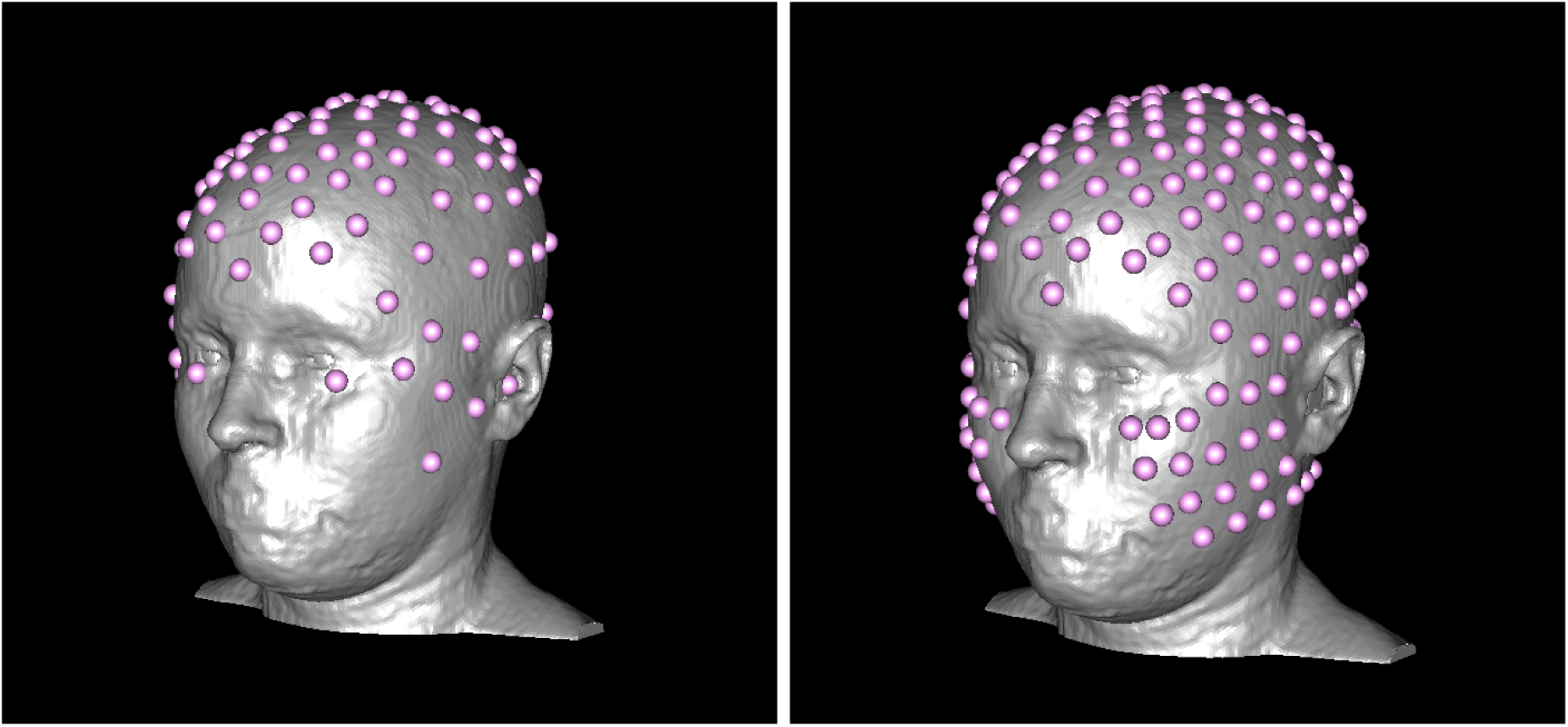
SCIRun manual registration of physical electrodes to head surface mesh: 128 electrodes *(left)*, 256 electrodes *(right)*.

The SCIRun networks for registration are included in the Appendix in Section A.5 in Figures 37 – 40.

### 2.3 MRI Segmentation of Tissues

We segmented the head volume using FSL and Seg3D, a free volume segmentation and processing tool,^17^ into air, cerebral spinal fluid (CSF), white matter, gray matter, skull, sinus, eyes, and scalp. We used FSL for initial segmentation and Seg3D for all further segmentation, including thresholding and manual editing.

We generated the initial brain segmentation with FSL from the *T*_1_ MRI by stripping the skull with the brain extraction tool (BET)^18^ (Figure 6) and then segmenting with FAST segmentation (Figure 7). FSL FAST outputs grayscale probability images of CSF, white matter, and gray matter layers (Figure 8) as well as a bias-corrected *T*_1_ MRI (Figure 9).

**Figure 6:**
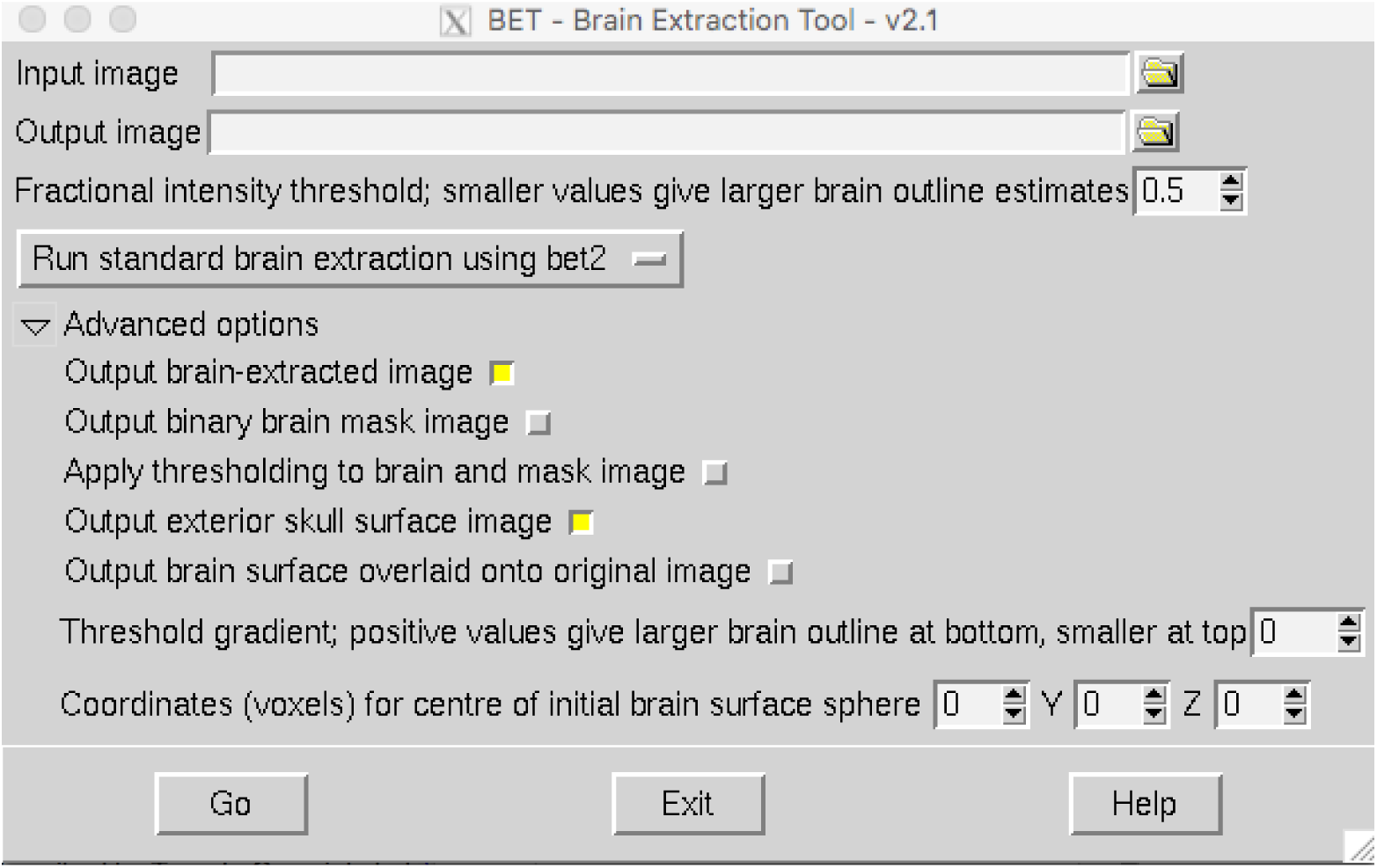
FSL’s BET2 tool.

**Figure 7:**
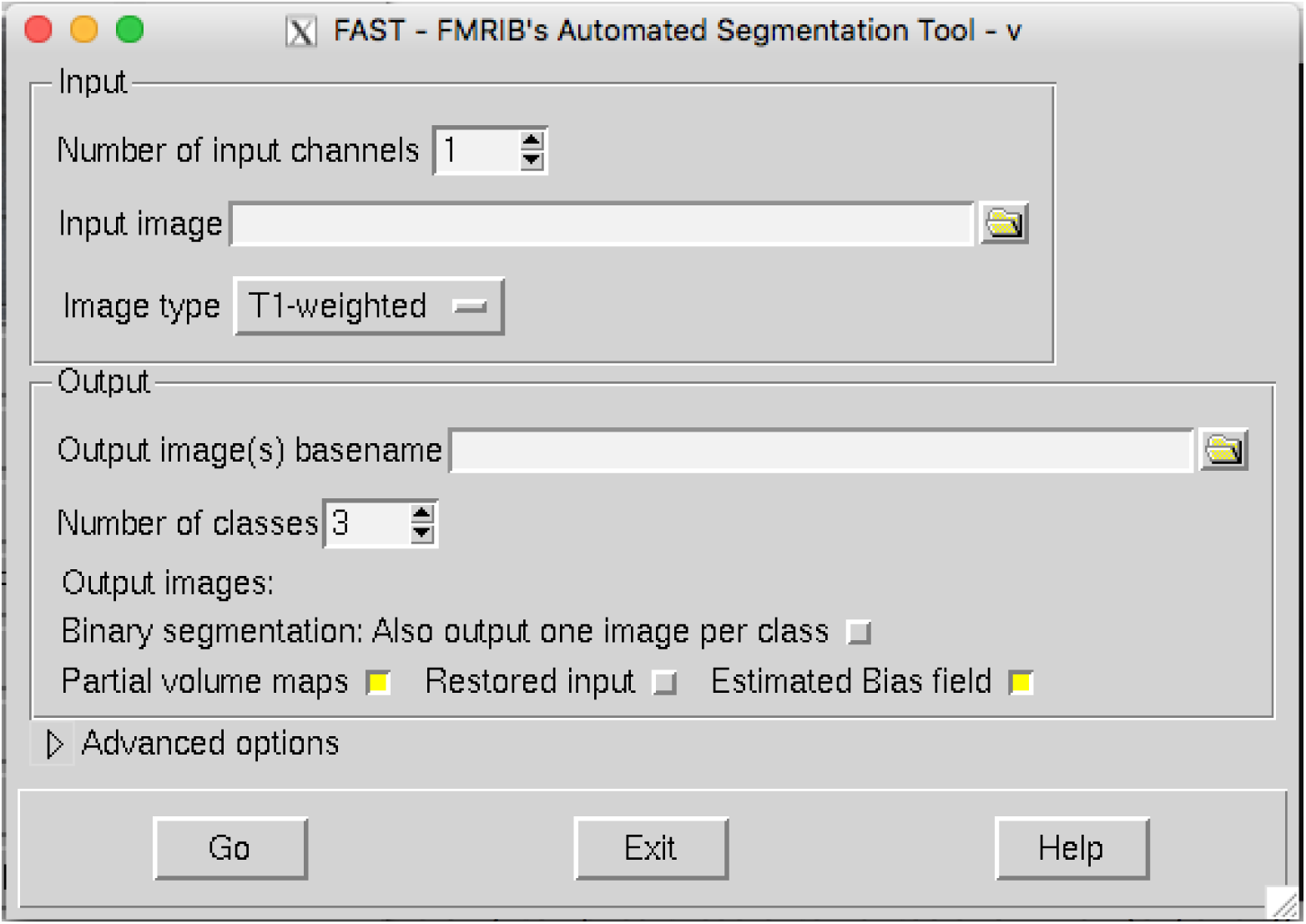
FSL FAST user interface.

**Figure 8:**
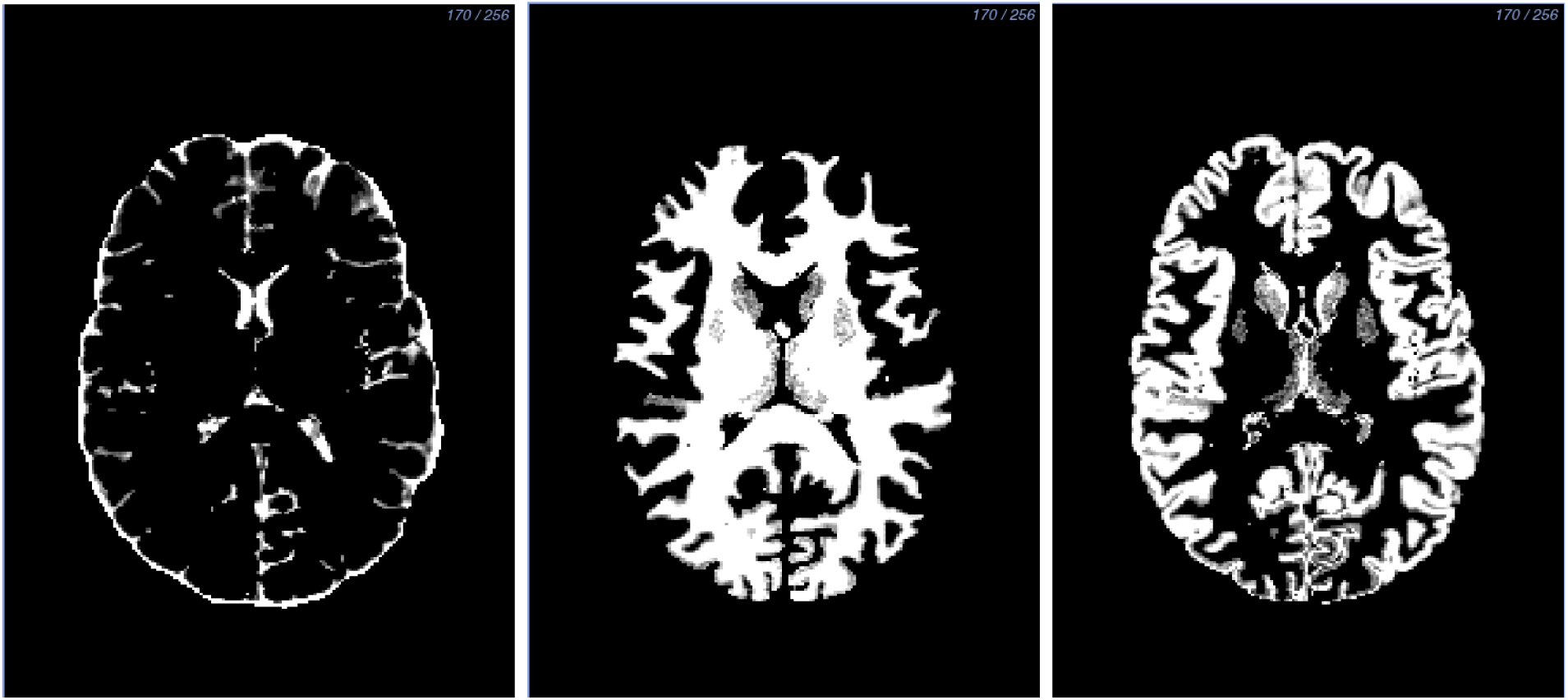
FSL FAST output: CSF *(left)*, white matter *(center)*, gray matter *(right)*.

**Figure 9:**
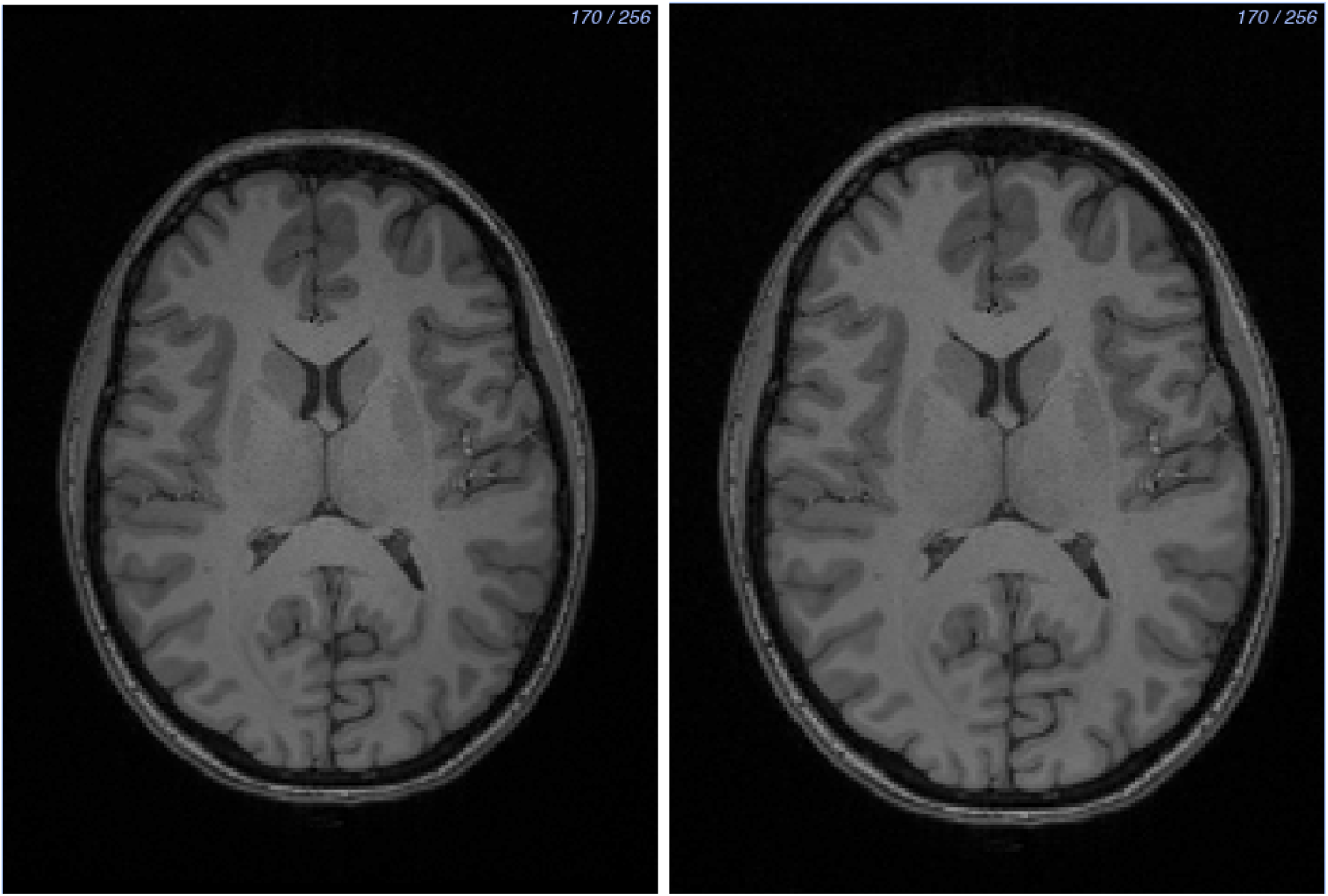
FSL FAST output: *T*_1_ MRI image before bias correction *(left)* and after bias correction *(right)*.

We created segmentations from the FSL FAST output by thresholding the probability images and by inspecting and manually editing each slice to remove any crossover between the layers, to add more detail, and to smooth out noisy areas. We started with the white matter layer because it is the innermost layer (Figure 10).

**Figure 10:**
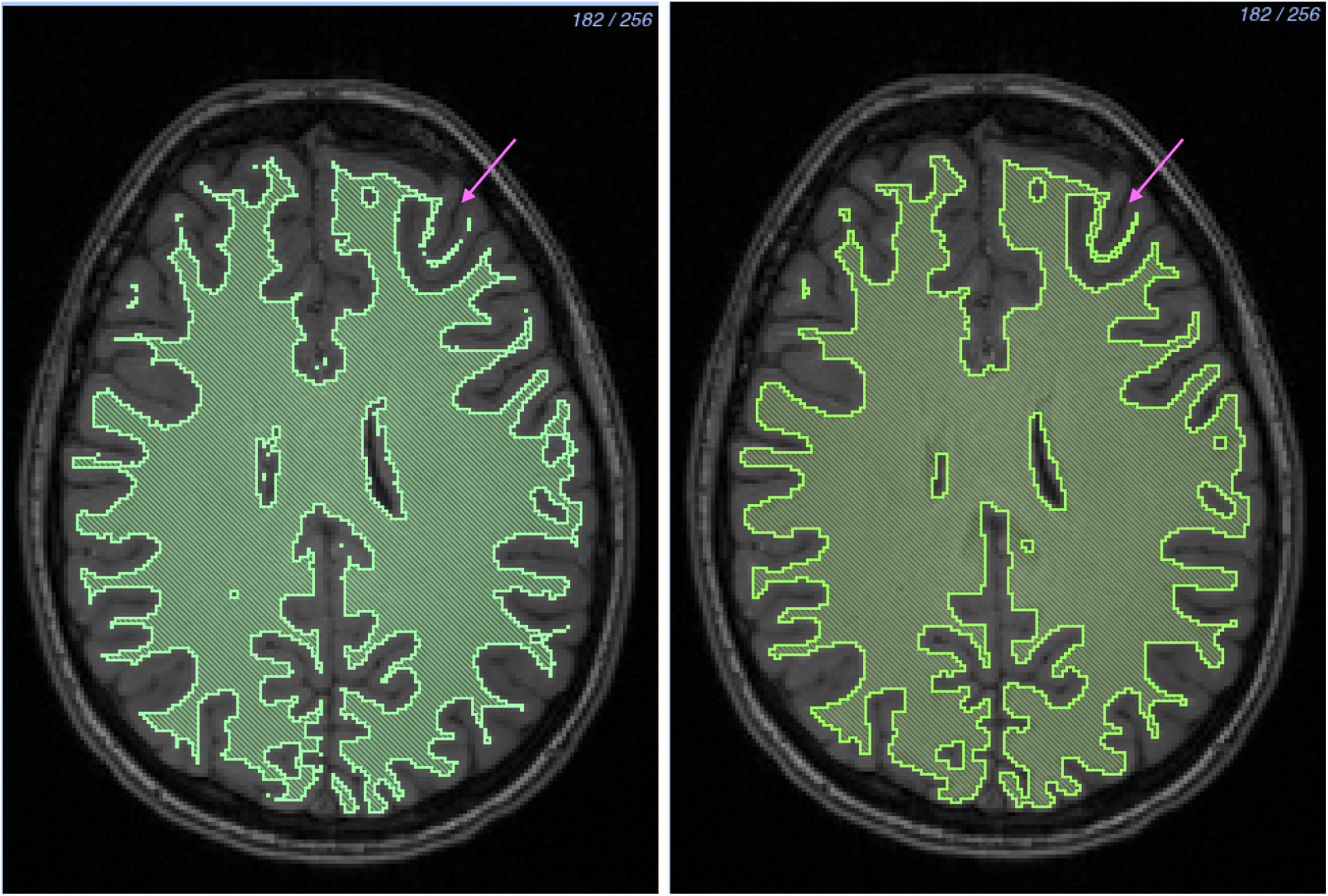
White matter segmentation: Before *(left)* and after *(right)* manual segmentation. The hook feature in the upper right-hand corner is a notable change between the two layers. The layer is more full and has less noise.

After we segmented the white matter, we created a threshold layer from the FSL FAST output for the gray matter. We then inspected and manually edited each slice in every direction. After manual editing, we removed the white matter layer from the gray matter using a Boolean remove mask filter to ensure no overlap between the layers. Lastly, we added a gray matter nuclei segmentation to the gray matter layer. The thresholding algorithms in Seg3D produced noise around these nuclei because of their grayscale similarities to white matter. To fix this noise, we segmented the nuclei area manually using the paintbrush tool in Seg3D. We added the nuclei to the gray matter layer using a Boolean OR mask filter, and removed any overlap from the white matter layer using a Boolean remove mask filter. We manually filled any holes between the two layers that could be assigned to either white or gray matter layers (Figure 11).

**Figure 11:**
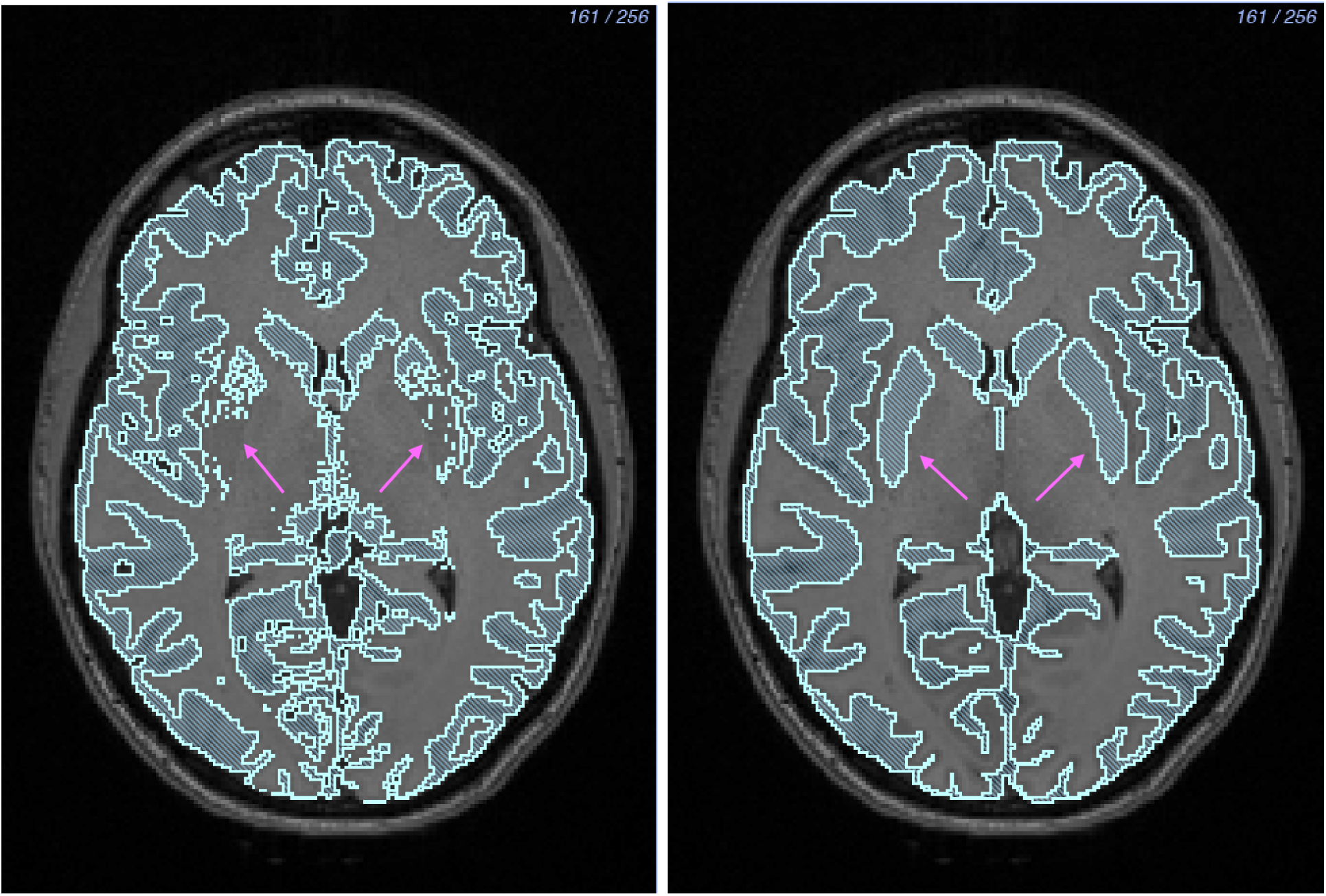
Gray matter segmentation: Before *(left)* and after *(right)* manual segmentation. Gray matter nuclei located in the center of the brain were segmented manually.

After we completed the segmentation of gray and white matter, we segmented CSF by creating a solid threshold layer for the entire brain and removing the white and gray matter layers using a Boolean remove mask filter (Figure 12). We then checked the white matter, gray matter, and CSF layers for holes, both on the surface and inside the segmentation between layers. We also performed a manual quality check on the layers to ensure that they were at least two pixels wide throughout. This thickness criterion helped to ensure a quality tetrahedral mesh.

**Figure 12:**
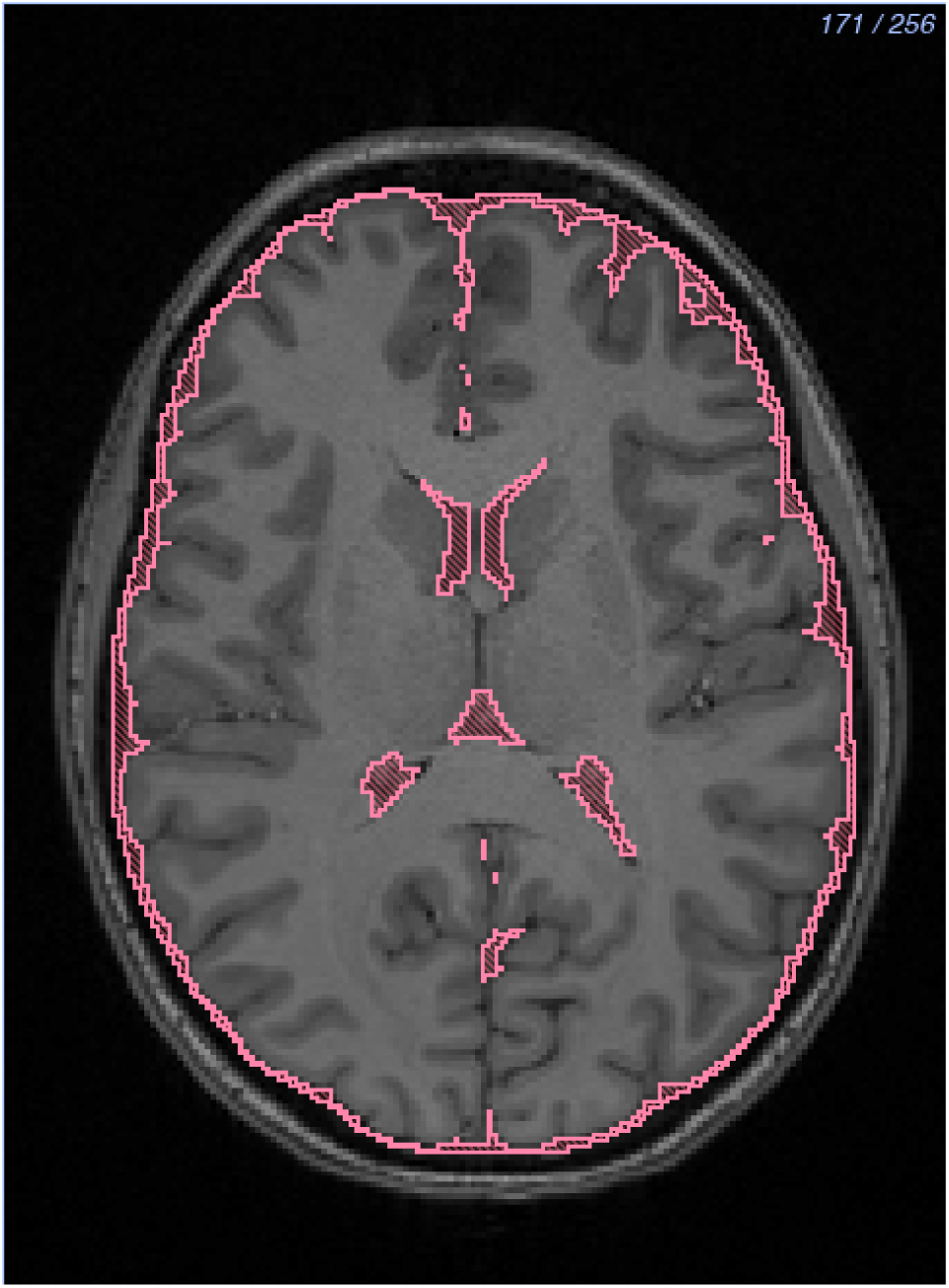
CSF segmentation.

We segmented the skull and sinus from an MR-based synthetic pseudo-CT image (Figure 20-*(left)*). We used an improved iterative version of the patch-based method, as described by Torrado-Carvajal et al.,^19^ that takes the *T*_1_ and *T*_2_ images as input and synthesizes the pseudo-CT based on both images, providing more refined and accurate bone boundaries.

For the skull segmentation, we applied a median filter with a one-pixel radius to the pseudo-CT image and thresholded the white pixels. We then manually edited each slice in every direction to add detail and to smooth noise (Figure 20-*(center)*). Since the subject had a permanent dental retainer that caused white pixels throughout her mouth in the pseudo-CT image, we segmented the mouth as solid bone. We deemed acceptable because the EEG cap used did not cover the subject’s mouth.

For the internal air segmentation, including the sinuses, esophagus, and ear canals, we thresholded the black pixels of the pseudo-CT image and manually edited each slice in every direction (Figure 20-*(right)*). We also performed a quality check on both layers to ensure they did not contain holes or have any layer overlap and were at least two pixels thick.

**Figure 13:**
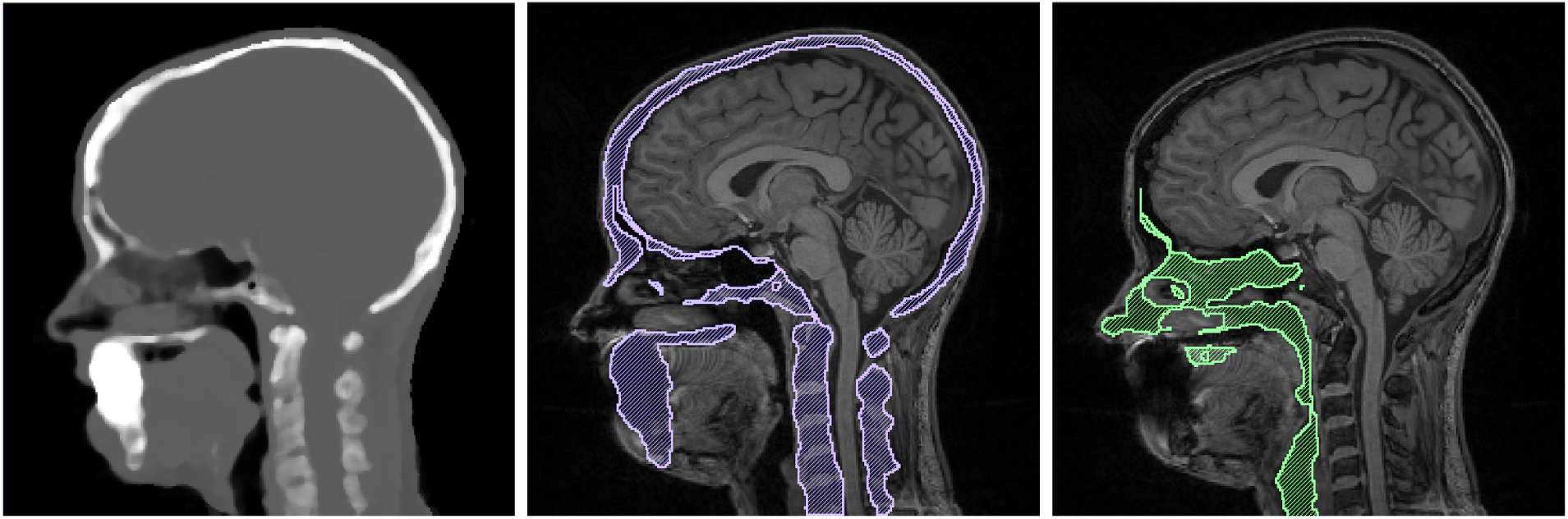
Skull segmentation: Pseudo-CT image *(left)*, skull layer *(center)*, and sinus layer *(right)*.

We then segmented the eyes, skin, and external air. We segmented the eyes by thresholding the *T*_2_ MRI, and the skin layer by thresholding the entire head volume and removing all previous segmentation layers using a Boolean remove mask filter. We performed a quality check on the skin layer to ensure that it was at least two pixels thick. The areas that required significant correction were the bridge of the nose, the bottom of the chin, and the sides of the head. Finally, we included pixels not previously assigned as the external air layer. We checked the entire segmentation to make sure there were no holes between layers to assure a quality mesh and accurate simulation results.

**Figure 14:**
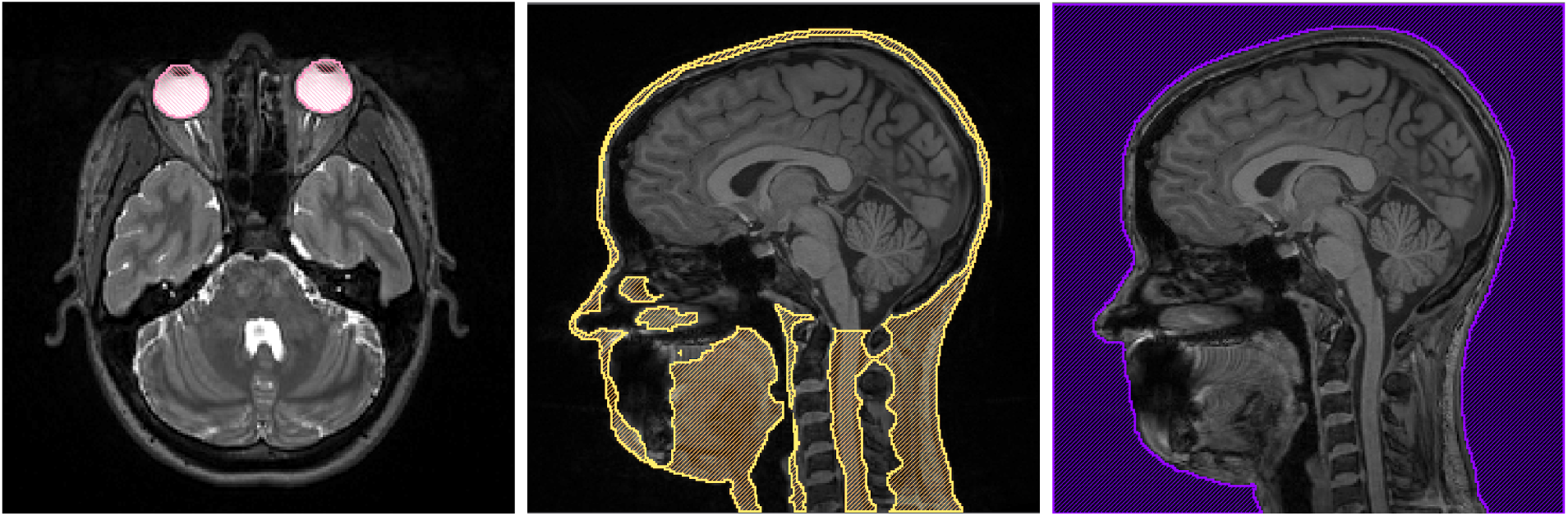
Eye segmentation with *T*_2_ MRI *(left)*, skin segmentation *(center)*, and air segmentation *(right)*.

The segmentation of the image data took approximately 100 hours, most of which was dedicated to manual editing. A breakdown of time per segmentation layer is shown in Table 1.

**Table 1:** Segmentation Time

| Segmented Tissue | Amount of Work (hrs) |
| --- | --- |
| White Matter | 40 |
| Gray Matter | 20 |
| CSF | 4 |
| Skull and Sinus | 35 |
| Eyes, Scalp, & Air | 8 |

### 2.4 Mesh Generation

We used our full-head segmentation to generate realistic three-dimensional geometries for use in subsequent finite element simulations. We generated a smooth, linear, subject-specific, boundary-conforming, quality, high-resolution, tetrahedral mesh using the Cleaver software^20^ on a Late 2013 Mac Pro with a 2.7 Ghz 12 Core Intel Xeon E5 processor with 64 GB of RAM and an AMD FirePro graphics card using the parameters listed in Table 2. Cleaver is a multimaterial meshing package that produces structured meshes of tetrahedral elements with guaranteed minimum element angles, resulting in quality meshes that require fewer computational resources.

**Table 2:** Clever Settings (High Resolution)

|  |  |
| --- | --- |
| scaling factor | 0.6 |
| size multiplier | 1.0 |
| lipschitz | 0.2 |
| padding | 0 |
| element sizing method | adaptive |

We created indicator functions, which describe the location of the surface with subgrid accuracy, by calculating inverted distance maps of each layer in the full-head segmentation in Seg3D using the distance map filter. To reduce the size of the mesh, we generated a new mesh, changing only the scaling factor parameter to 1.0 from the parameters in Table 2. We exported the computing sizing field from Cleaver and manipulated it in SCIRun by changing how quickly the elements increased in size. We input the changed sizing field into Cleaver with the same indicator functions and successfully cleaved a new, smaller, quality mesh. Both sizing fields are included in the dataset.

### 2.5 Mathematical Modeling

The forward and inverse EEG problems are governed by a generalized Poisson equation (1). We used the head mesh, with associated inhomogeneous and anisotropic conductivity regions, as a volume conductor to solve the following boundary value problem:

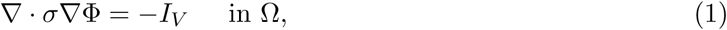

where Φ is the electrostatic potential, *σ* is the electrical conductivity tensor, and *I_V_* is the current per unit volume defined within the solution domain, Ω. For the forward EEG problem, we solved equation 1 for Φ with a known description of *I_V_* and the Neumann boundary condition:

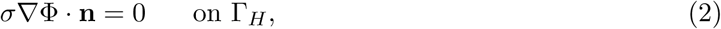

which says that the normal component of the electric field is zero on the surface interfacing with air (here denoted by Γ_*H*_). For testing purposes, we used dipoles for the current source. We calculated the electrical and potential fields everywhere within the head model.^2^

#### 2.5.1 Electrical Conductivity Preparation

All electrical conductivities were homogeneous for each tissue with the exception of the white matter when using DTI data. The isotropic conductivities^21^ we used are shown in Table 3.

**Table 3:** Isotropic Tissue Conductivity

| Tissue Type | Isotropic Conductivity ($S/m$) |
| --- | --- |
| White Matter | 0.1429 |
| Gray Matter | 0.3333 |
| Cerebrospinal Fluid (CSF) | 1.79 |
| Skull | 0.001 |
| Skin | 0.4346 |
| Sinus | 1e-6 |
| Eyes | 0.5051 |

When we added the DTI tensor data, we used two approaches to convert the tensor data to conductivities. The first was scaling the data:^22^

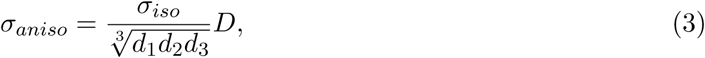

where *D* is the diffusion data, *d_i_* is the *i*th eigenvalue of *D*, and *σ_iso_* is the white matter isotropic conductivity. The second method gave the white matter a fixed ratio of conductivity:

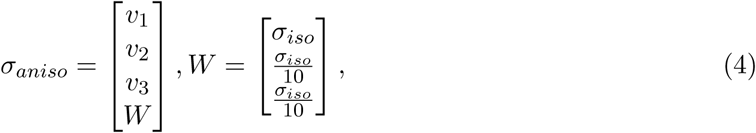

where *v_i_* is the *i*th eigenvector of *D*, *W* is the white matter ratio vector, and the ratio is 10 : 1.

We implemented both methods into the SCIRun networks for anisotropic forward problems (Figure 38).

### 2.5.2 Numerical Methods

We computed solutions to equation 1 using the finite element method. By applying Green’s theorem to equation 1, we generated the following weak formulation:

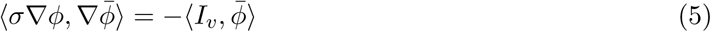

where 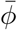 is an arbitrary test function, which can be thought of physically as a virtual potential field. By applying the Galerkin approximation to equation 5, we can represent the finite element approximation as

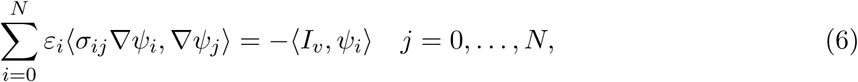

subject to the Dirichlet boundary condition. The finite element approximation of equation 1 can equivalently be expressed as a system of *N* equations with *N* unknowns *ε_i_ … ε_N_* (e.g., the electrostatic potentials). In matrix form, the above system can be written as *Aε* = *b*, where *A* = (*a_ij_*) is called the global stiffness matrix and has elements (*a_ij_*) = (*σ_ij_∇ψ_i_, ∇ψ_j_*), and *b_i_* = *−*(*I_v_, ψ_i_*) is usually termed the load vector. For volume conductor problems, *A* contains all the geometry and conductivity information of the model.^2^

We used SCIRun to apply parameters and to solve equation 6 numerically using linear basis functions for tetrahedral elements. Within the SCIRun environment, we applied isotropic and anisotropic conductivity tensors to the tetrahedral mesh as well as to inhomogeneous regions. We applied Dirichlet and Neumann boundary conditions to compute potentials using a conjugate gradient method with a Jacobi preconditioner (Figures 37 – 38).

### 2.6 Simulations and Visualizations

We performed all simulations and visualizations in SCIRun. All networks are shown in the Appendix in Section A.5 – Figures 36 – 40.

#### 2.6.1 Forward Problem

Solving systems from a known source to the EEG electrodes, described in Section 2.5, is known as a forward problem. The opposite action, solving systems from the EEG data to a unknown source, is an inverse problem. In this project, we built SCIRun example networks to solve forward problems with known sources and to write a lead field matrix for use in future inverse problems. We solved forward problems with isotropic and anisotropic conductivities, using DTI data for the direction of anisotropic conductivities.

The required inputs for the isotropic example network were the tetrahedral head mesh, isotropic conductivities, the head segmentation, the physical electrode locations, and dipole sources. The SCIRun network allowed the user to choose a dipole as a current source from the set for a forward simulation. The network also removed fiducial electrodes from the dataset. We registered the mesh to the head segmentation using the rigid registration previously described in Section 2.2.6. We then registered the electrodes and the dipoles to the mesh coordinate space, as described in Section 2.2.6. After we removed any flat tetrahedra from the mesh, we mapped the conductivities to their respective tissues. We solved the system with the mapped data and the chosen dipole sources. We then extracted the solution onto the head surface and the electrode locations for visualization. We also used streamlines and isopotential lines for visualization.

For the anisotropic example, the network was similar to the isotropic example with the exception of the scaled diffusion tensor data used as conductivities for white matter, and the DTI to mesh transform as additional input. We registered the head mesh, electrodes, and dipoles to the DTI space with a rigid registration, as described in Section 2.2.6; the head segmentation was not needed.

#### 2.6.2 fMRI

We visualized the fMRI data one time step at a time using the two-dimensional matrix, described in Section 2.2.4, as input by extracting one column at a time. We registered the fMRI to the tetrahedral mesh as described in Section 2.2.6. After registration, we mapped the smoothed fMRI data onto the mesh using a mapping matrix with a linear interpolation basis.

#### 2.6.3 EEG

We visualized EEG data on the physical electrode locations, which we registered to the head mesh with a rigid registration, as described in Section 2.2.6, after removing the fiducials from the electrode dataset. We obtained the filtered EEG data one time step at a time by extracting the column of the matrix. We then placed the electrodes onto the mesh and mapped the EEG data onto the electrodes. An example EEG network is shown in Figure 40. Each EEG electrode configuration has its own SCIRun network.

## 3 Results & Discussion

The open-source dataset that was released with this paper includes raw image data, image acquisitions, and preprocessed/corrected images for all image modalities along with the necessary inputs and outputs for the preprocessing/correction steps of the pipeline. We also include the raw and preprocessed EEG signal data along with the scripts used to obtain and process the data. We include all intermediate steps in the dataset as well, such as the complete, high-resolution brain segmentation that can be used to create three-dimensional tetrahedral volume and surface meshes. We provide two three-dimensional tetrahedral finite element meshes made from the segmentation of different resolutions that serve as volume conductors to solve forward and inverse EEG problems along with the sizing fields used to generate them. We also provide simulation examples of the EEG forward problem, with isotropic and anisotropic systems; functional image data mapped onto a tetrahedral mesh; electroencephalography (EEG) signals mapped onto net electrodes; and diffusion tensor data.

All SCIRun networks used to generate results and the open-source dataset are available at www.sci.utah.edu/SCI_headmodel.

### 3.1 Diffusion Tensor Images

To add anisotropy to the forward problem simulations, which creates more accurate results, we generated a tensor field of the white matter tracts. Figure 15 and Figure 16 show the anisotropy in a typical axial cross section slice calculated from the preprocessed DWIs using equation 3. The anisotropy of the white matter is represented by ellipsoids (Figure 16). The direction and radius of the ellipsoid correspond to the eigenvector and eigenvalue of the anisotropy tensor for each voxel, respectively. The RBG color maps to the XYZ direction of the primary eigenvector.

**Figure 15:**
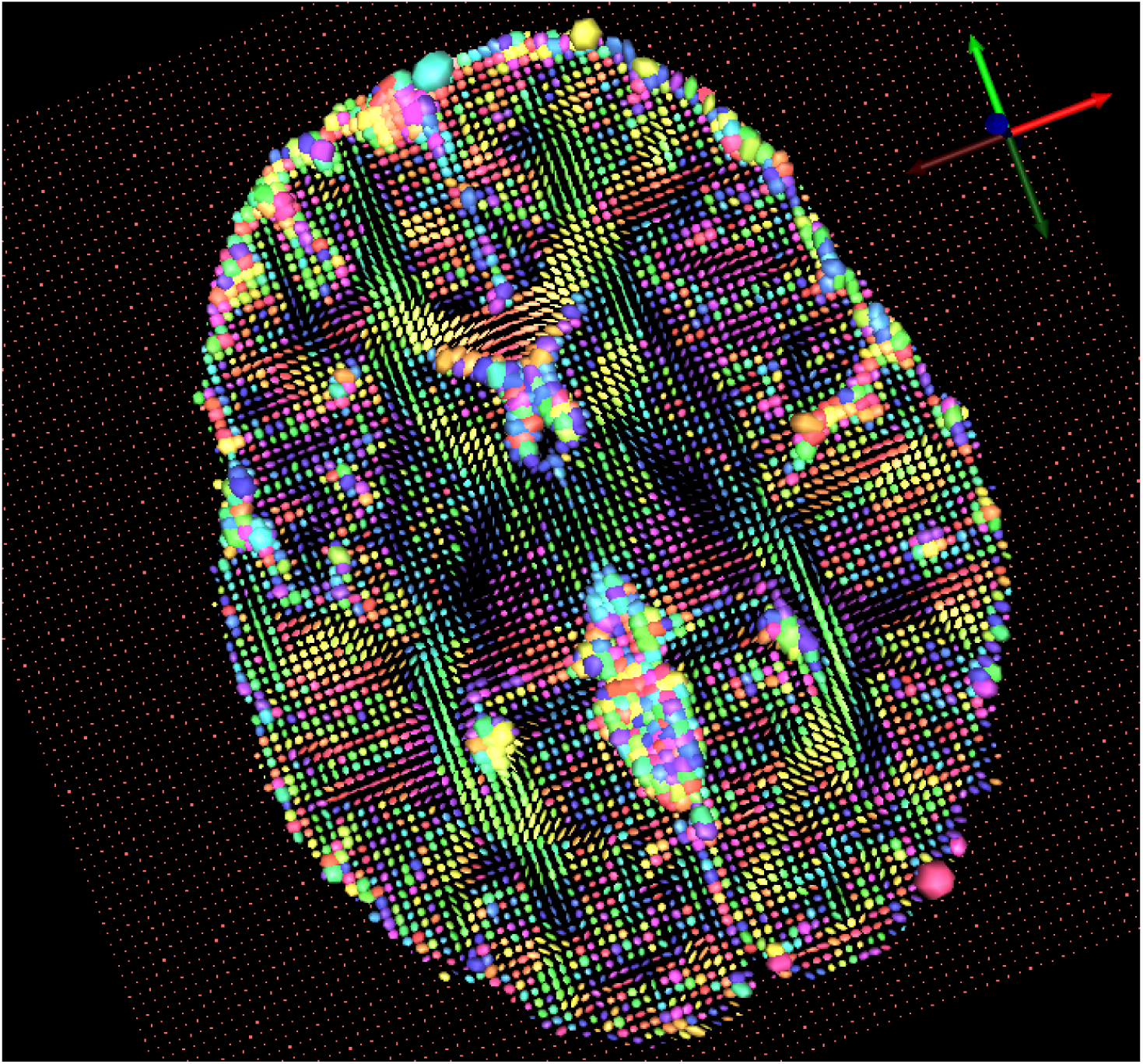
Diffusion tensor visualization using SCIRun: a typical axial slice.

**Figure 16:**
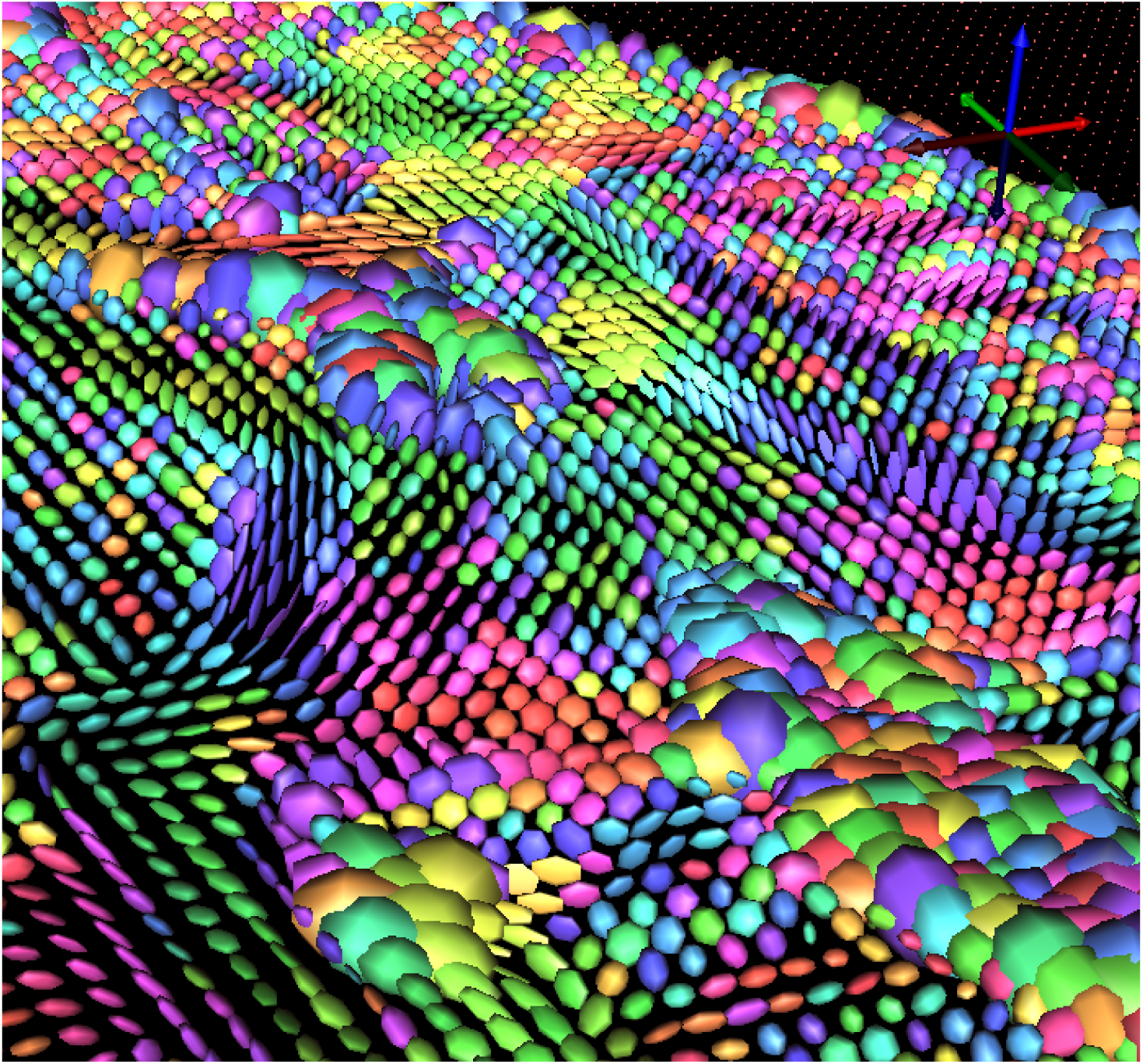
Diffusion tensor visualization using SCIRun: anisotropy represented by ellipsoids.

The manual registration we applied to the diffusion tensor data, described in Section 2.2.6, aligned well with the cortical mesh (Figure 4 (*left*)). Figure 15 shows more anisotropy throughout the white matter and less throughout the gray matter and CSF. However, the alignment was imperfect resulting in noise around the outside edge of the axial slice.

We chose to build the tensor field in SCIRun rather than in 3D Slicer^23^ or FSL DTIFIT, which are common software packages for building brain tensor fields, because the orientation of data from these packages differed in orientation when compared to SCIRun. The difference in orientation made for difficult registration to the cortical surface mesh, which we performed using SCIRun. Tensor orientation is important when registering to diffusion data and can be difficult to correctly register. We registered other datatypes to the diffusion tensor coordinate space to ensure the tensor orientation was correct. When we built the tensor field in SCIRun and put it back into 3D Slicer, the data’s orientation appeared upside down and backwards (Figure 17) from what we saw in SCIRun. To apply a correct and accurate registration, we used the outputs of DTIFIT in SCIRun as explained in Section 2.2.6.

**Figure 17:**
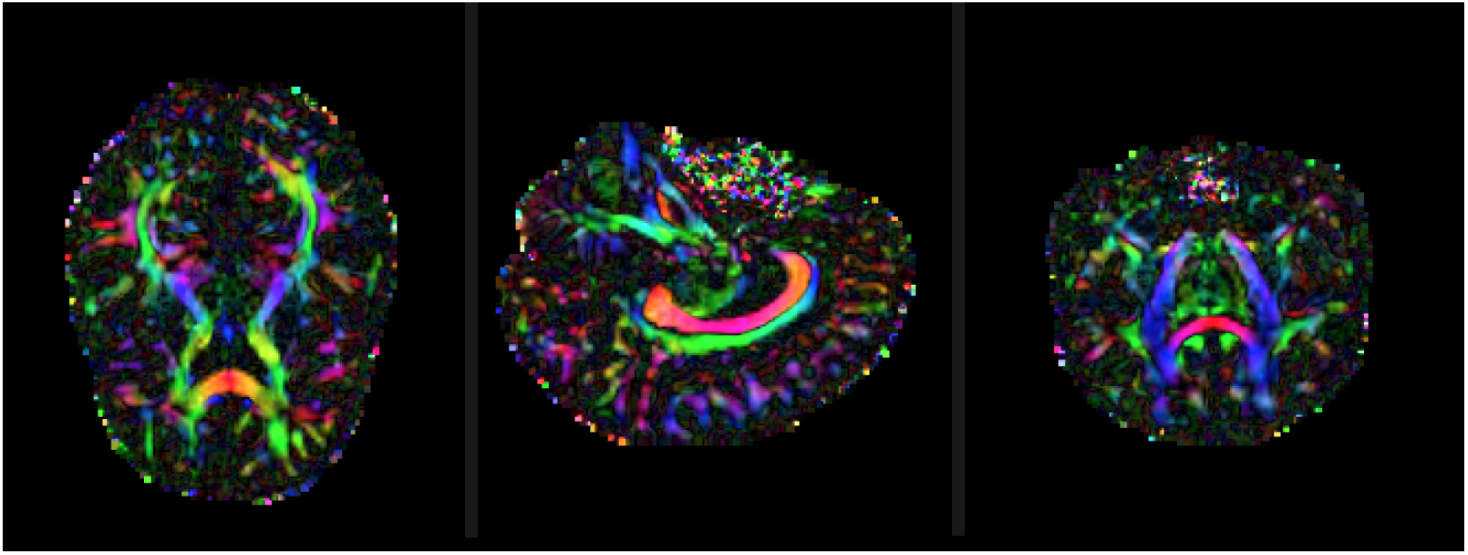
Example of difference in orientation between SCIRun and 3D Slicer data.

### 3.2 Segmentation

For this project, we segmented the head into eight detailed layers (Figure 18), listed in Section 2.3. We used this segmentation to create an inhomogeneous three-dimensional tetrahedral mesh.

**Figure 18:**
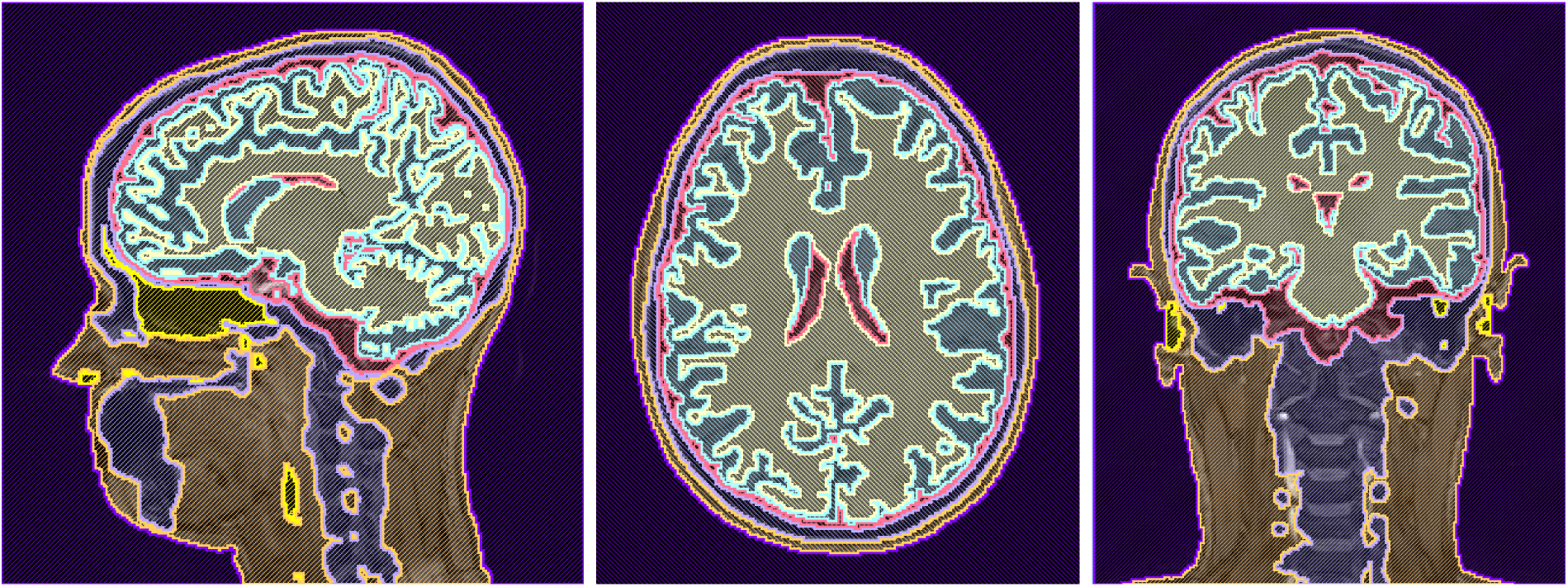
A high-resolution, eight-layer, full-head segmentation made with Seg3D.

Segmentation of the brain was difficult because of the similar grayscale intensities across different tissues, especially between white and gray matter; thresholding the image produced noisy and incomplete layers (Figure 19). As discussed in Section 2.3, we used FSL FAST and Seg3D for segmentation. This method, compared with Freesurfer,^24^ Statistical Parametric Mapping through Matlab (SPM),^25^ Atlas Based Classification through 3D Slicer,^26^ and Seg3D methods alone, produced the most qualitatively accurate head and brain segmentation results for this data.

**Figure 19:**
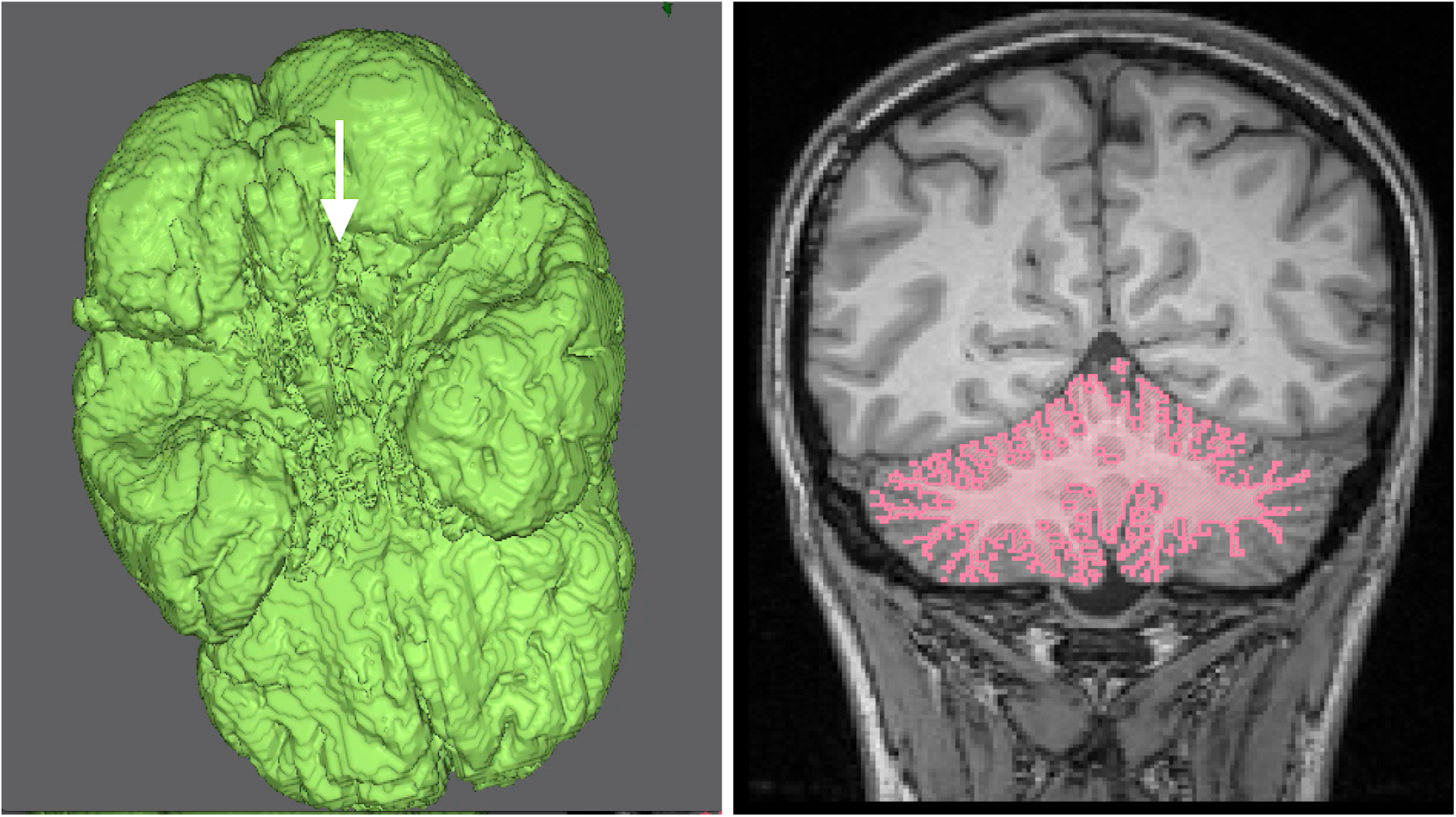
Noisy and incomplete segmentation layers: Isosurface segmentation of gray matter *(left)* and coronal slice of cerebellum segmentation *(right)*. Both segmentations were made using Seg3D.

The skull and the sinus layers were the most difficult to segment using only an MRI, because they are represented by only black pixels with no clear tissue boundaries, and the subject’s data did not include a computed tomography (CT) scan. For our first attempt to create a bone layer, we used FSL’s BET2 tool (Figure 6) to extract a skull surface. We then thresholded the *T*_1_ MRI to create the remainder of the bones in Seg3D, and connected the bones to the skull made from FSL using a Boolean OR mask filter. Although this approach gave an adequate segmentation for the skull (Figure 20 – *left*), it did not include important tissues, such as the sinus layer.

This method, compared to the method described in Section 2.3, produced geometries that were similar in some areas but different in others. The segmentation created using FSL’s BET2 and Seg3D was rough and had a clear line of where the two segmentations were connected. It also did not include a chin or a sinus layer. The segmentation created from the pseudo-CT image was smooth and included a chin (Figure 20 – *right*) and a sinus layer (Figure 21). However, this segmentation included the inside of the mouth as bone, but as mentioned in Section 2.3, this inclusion was not a concern for simulations results.

**Figure 20:**
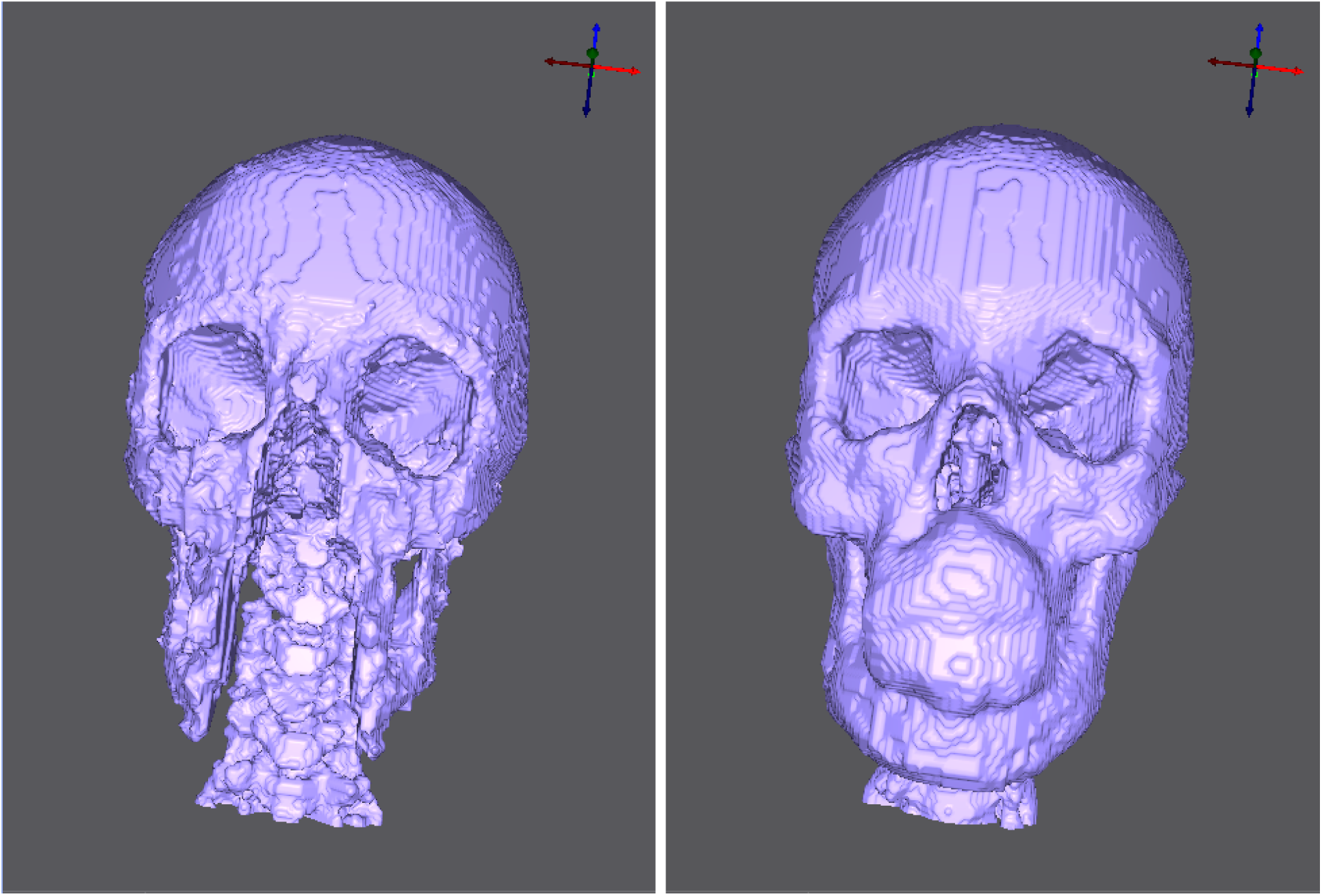
Skull segmentation comparison: Created with BET2 and thresholding *(left)* and with pseudo-CT image *(right)*. Both segmentations were made using Seg3D.

**Figure 21:**
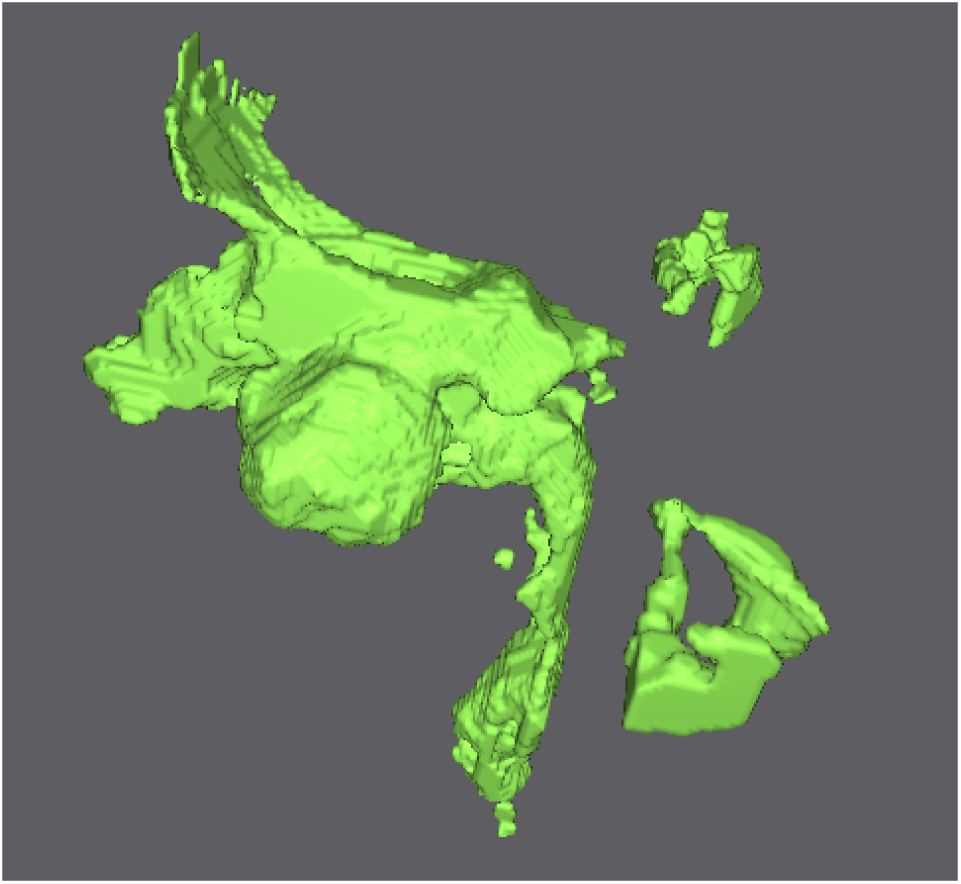
Sinus segmentation.

During MRI imaging, the subject was on her back, which caused the brain to shift slightly to the back of the head, causing thin segmented regions on the back of the head. Other thin segmented regions were on the side of the subject’s head, the bridge of the nose, and the bottom of the chin (Figure 22). We made these sections at least two pixels thick to ensure a quality mesh.

**Figure 22:**
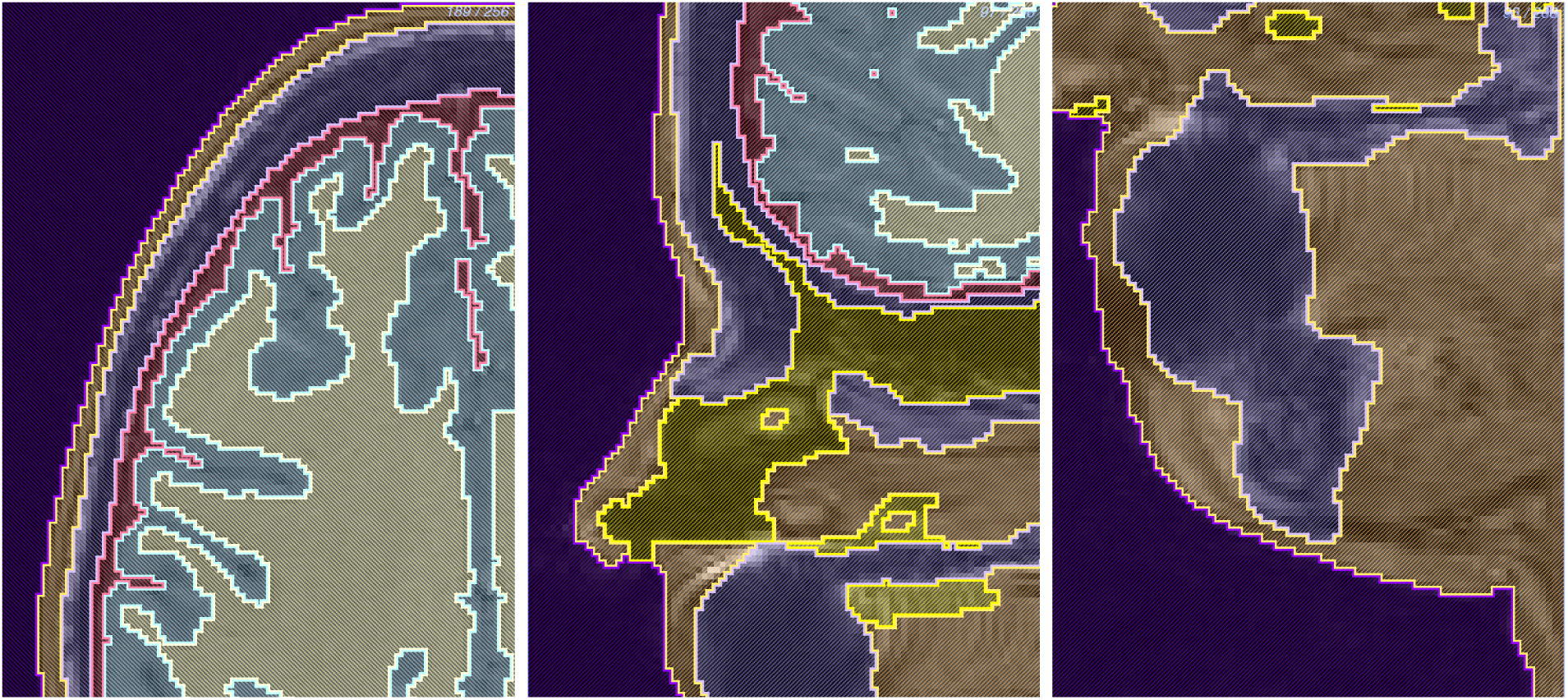
Thin segmentation regions: side of the head *(left)*, bridge of the nose *(middle)*, bottom of the chin *(right)*.

### 3.3 Finite Element Meshes

The highest resolution mesh we generated with the settings listed in Section 2.4 contained 60.2 million elements and 10.3 million nodes (Figure 23). This mesh was large because of the complexity of the segmentation, including small features, thin sections, and the multimatieral interface with three or more layers interacting at once. The simulations performed slowly when using this mesh because of it’s large size and required at least 32GB of RAM.

**Figure 23:**
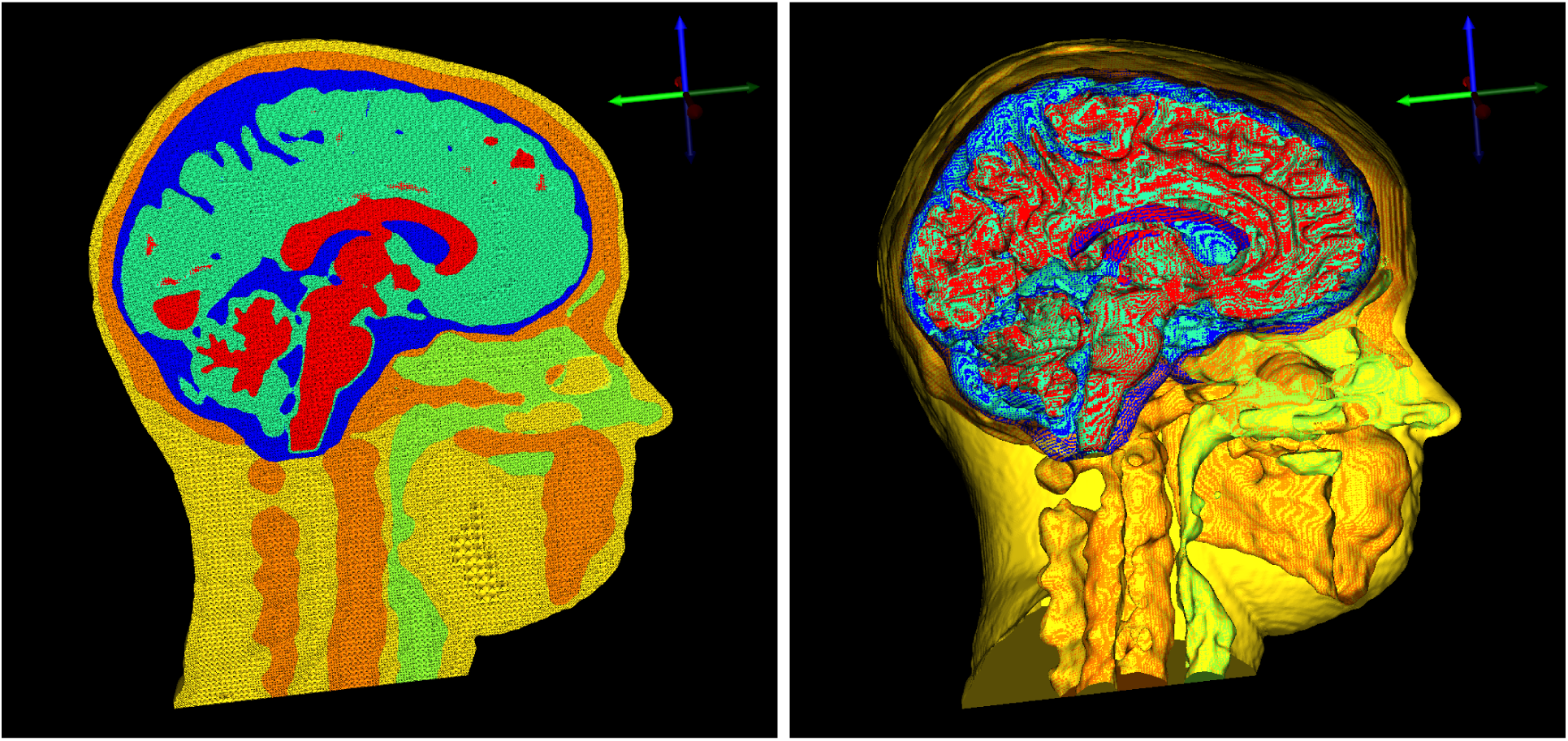
60.2 M element mesh: tetrahedral mesh *(left)*, surface mesh *(right)*.

We attempted to generate smaller meshes to be able to run faster simulations, but many of the meshes contained holes (Figure 24). After we manually changed the sizing field described in Section 2.4, we generated a mesh with 15.7 million elements and 2.7 million nodes without holes (Figure 25). However, this mesh contained one flat tetrahedra, which we later removed in a SCIRun network (Figure 37 and Figure 38). The production of flat tetrahedra in Cleaver is currently being investigated by Cleaver software developers.

**Figure 24:**
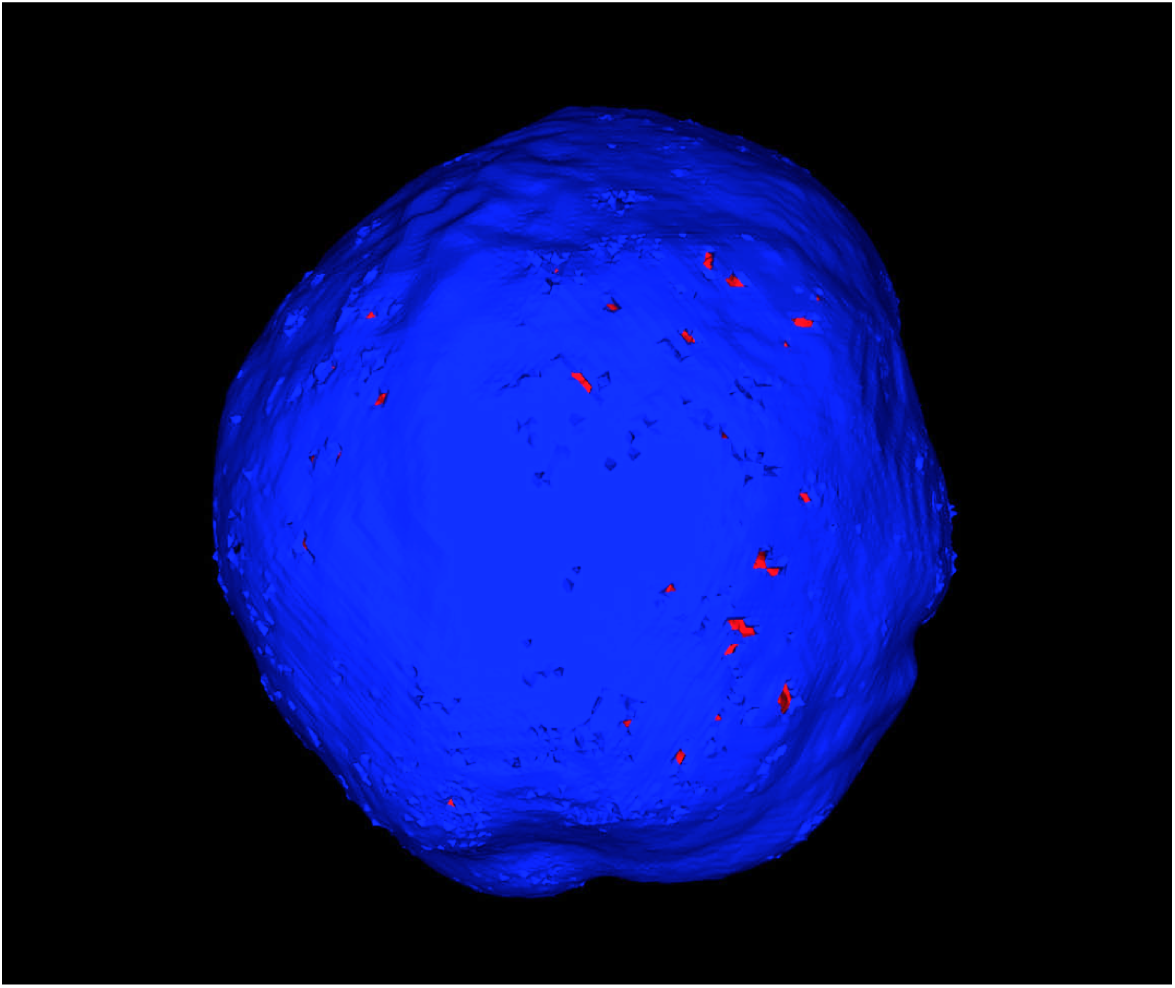
An example of a mesh containing holes.

**Figure 25:**
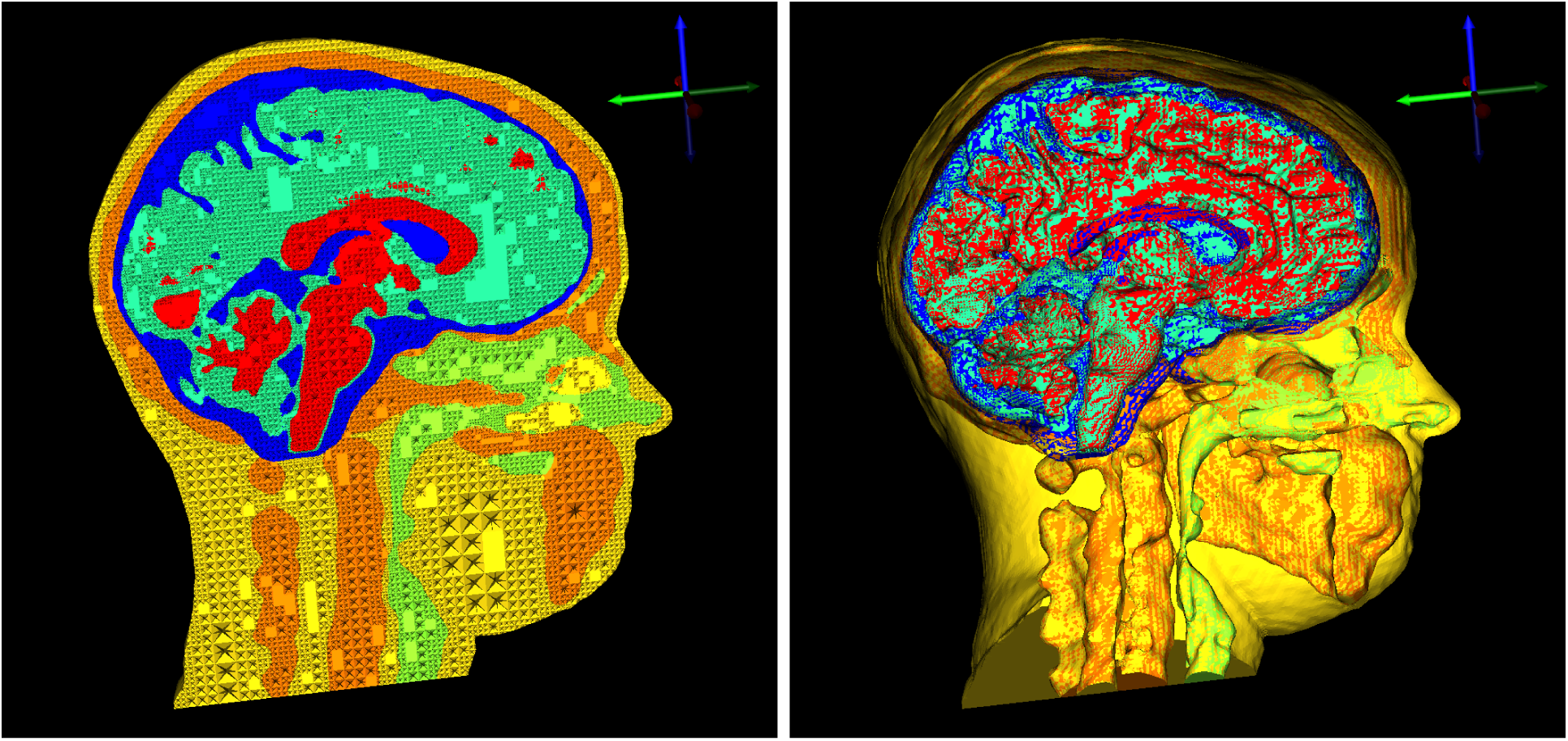
15.7 M element mesh: tetrahedral mesh *(left)*, surface mesh *(right)*.

### 3.4 Forward Problem

#### 3.4.1 Isotropic

An isotropic, inhomogeneous head model is expected to have mostly spherical propagation of electrical signals. The simulations showed spherical propagation and acceptable registration of electrodes and dipoles to the mesh space (Figure 26). We generated three-dimensional streamlines (Figure 27) as well as isopotential lines to visualize this propagation and to compare isotropic and anisotropic conductivity (Figure 30).

**Figure 26:**
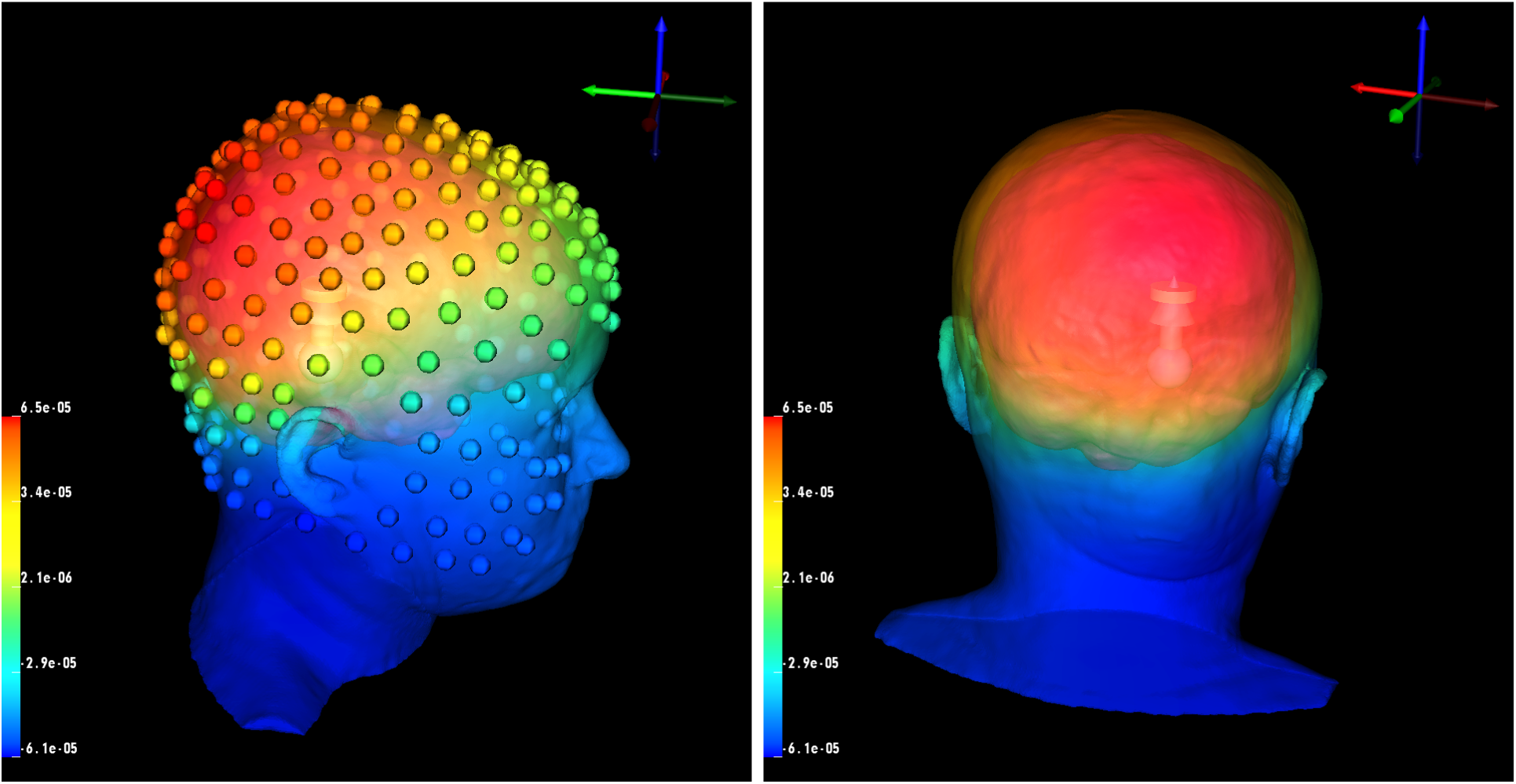
Isotropic forward problem solution with a dipole source and data mapped onto the head surface and electrodes.

**Figure 27:**
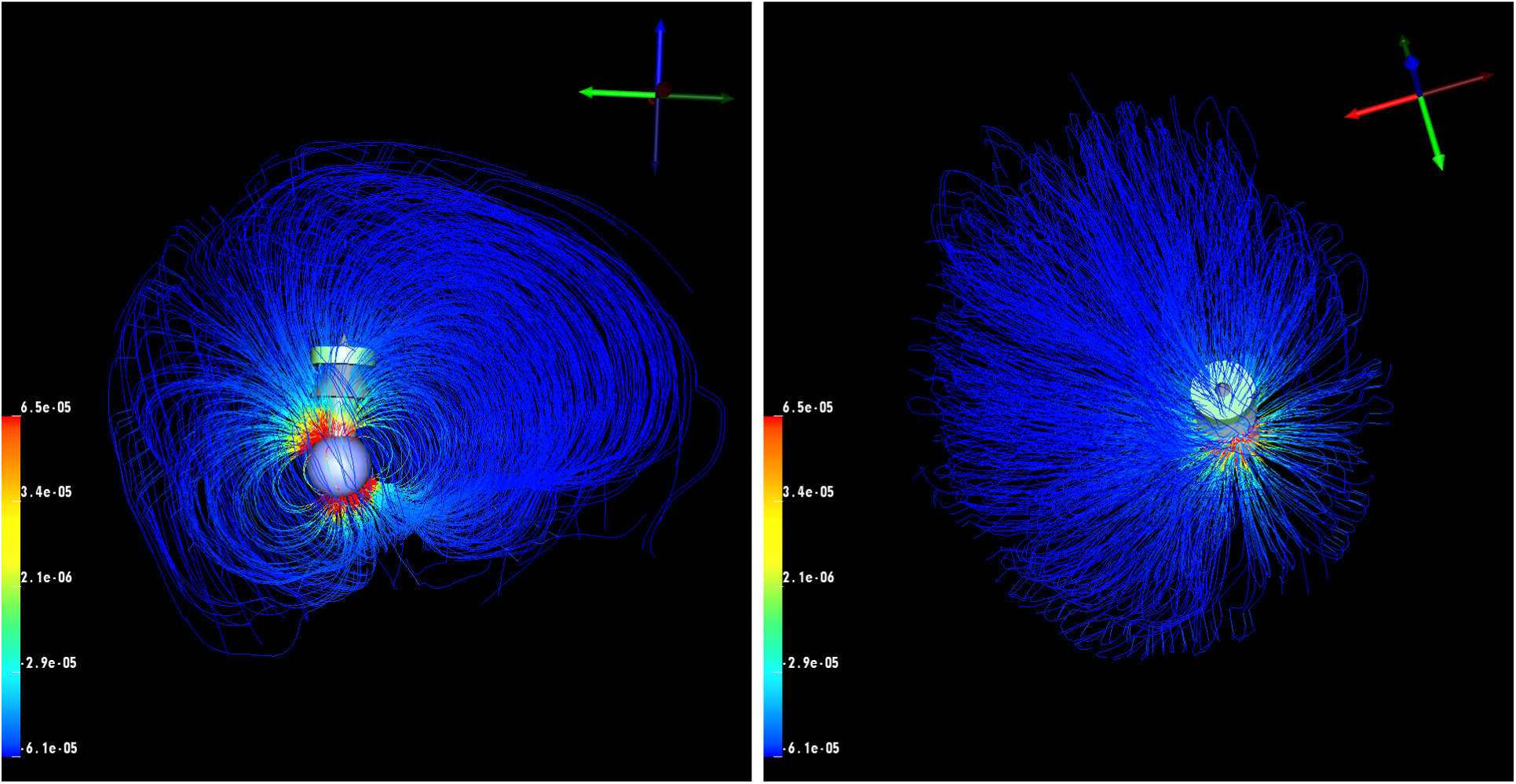
Isotropic streamlines with a dipole source.

#### 3.4.2 Anisotropic

The expectation for an anisotropic, inhomogeneous head model is to have nonspherical propagation, which can be seen with the streamline (Figure 29) and isopotential line (Figure 30) visualizations. As discussed in Section 2.5.1, two methods are used for implementing diffusion tensor data, scaling and using a fixed ratio, both available in the SCIRun network shown in Figure 38. For Figures 28 – 30, we used the scaling method (equation 3). The simulations showed nonspherical propagation and acceptable registration of the electrodes, dipoles, and mesh into the diffusion tensor space (Figure 28).

**Figure 28:**
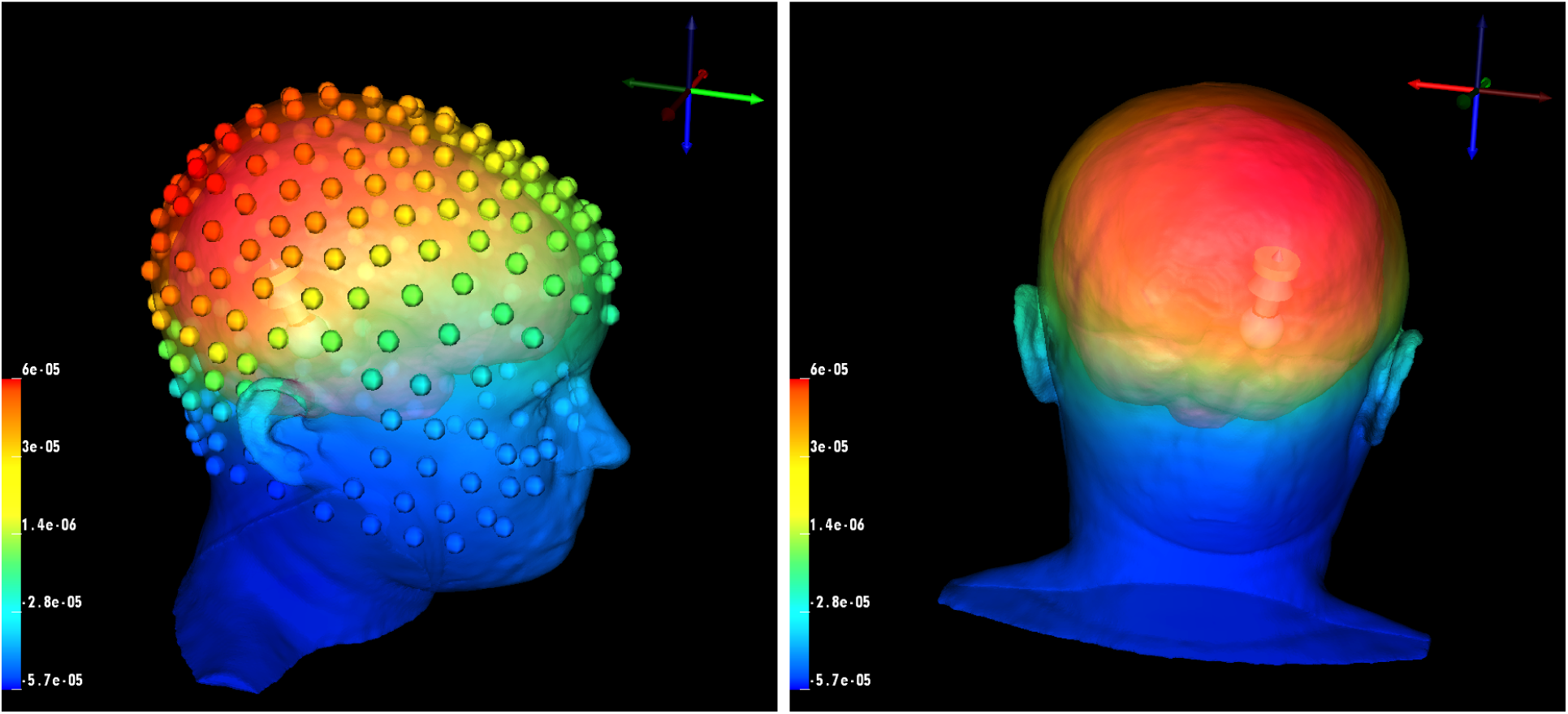
Anisotropic forward problem solution with a dipole source and data mapped onto the head surface and electrodes.

**Figure 29:**
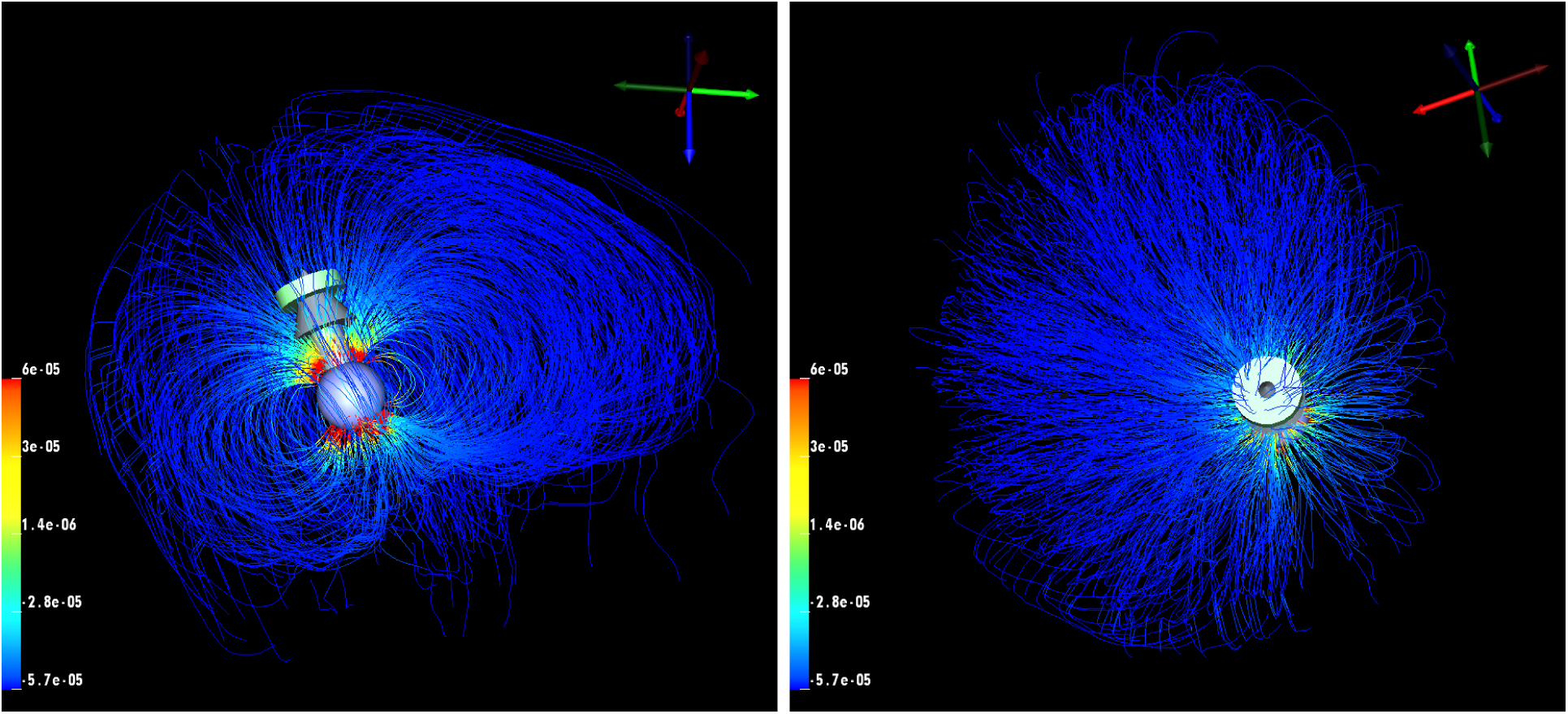
Anisotropic streamlines visualization with a dipole source.

**Figure 30:**
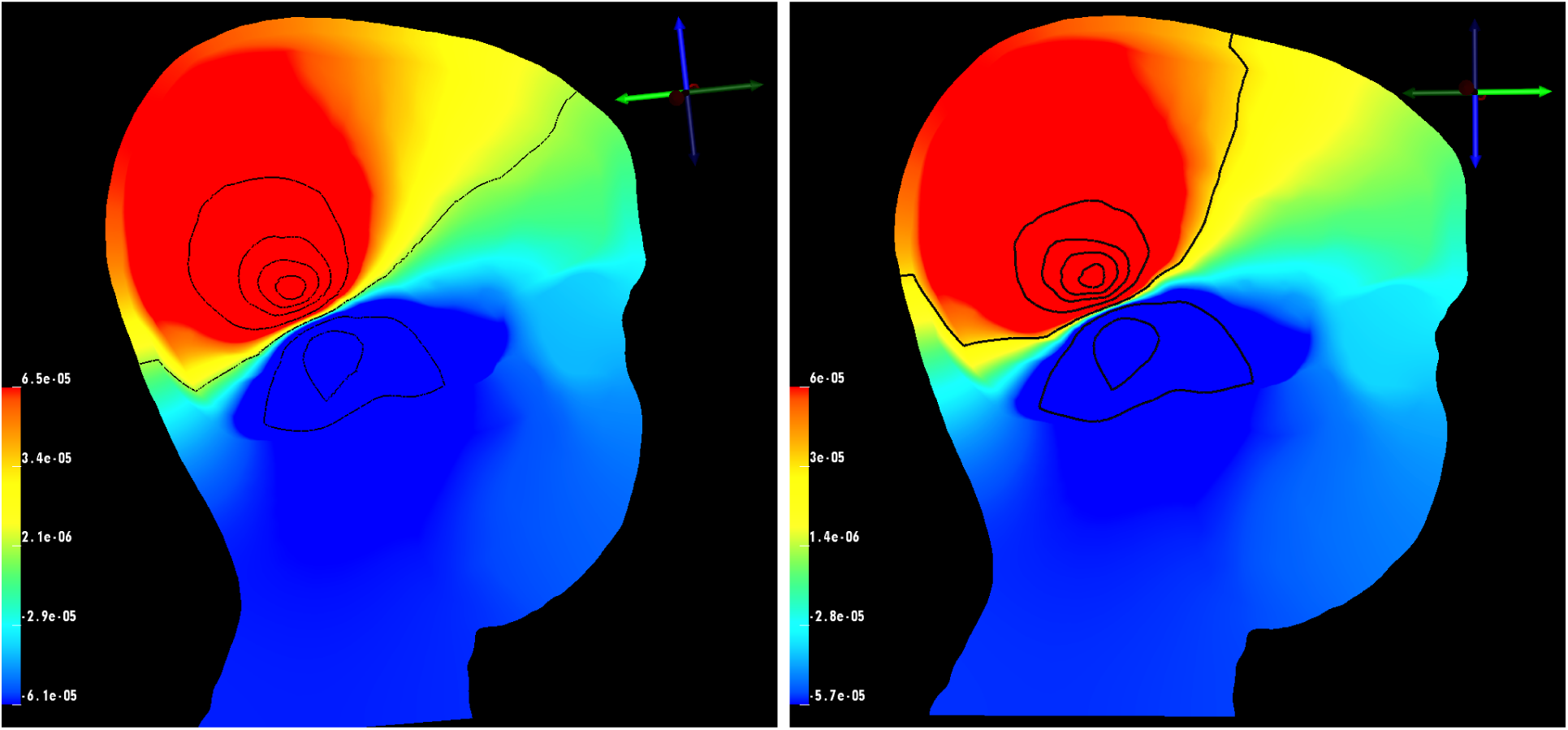
Isopotential lines comparison: isotropic white matter conductivity *(left)*, anisotropic white matter conductivity *(right)*.

### 3.5 fMRI Visualization

fMRI data was a novel imaging datatype for SCIRun. We successfully mapped and visualized the fMRI data onto the cortical surface with a rigid and manual registration, as discussed in Section 2.2.6, to the mesh coordinate space (Figure 31). This mapping network allows for future use of fMRI data in simulations using SCIRun (Figure 39). The manual registration we applied is fairly accurate since we did not see any large patches of zero signal, which would appear in blue. We also saw the higher signals on the brain stem, which has the most blood flow.

**Figure 31:**
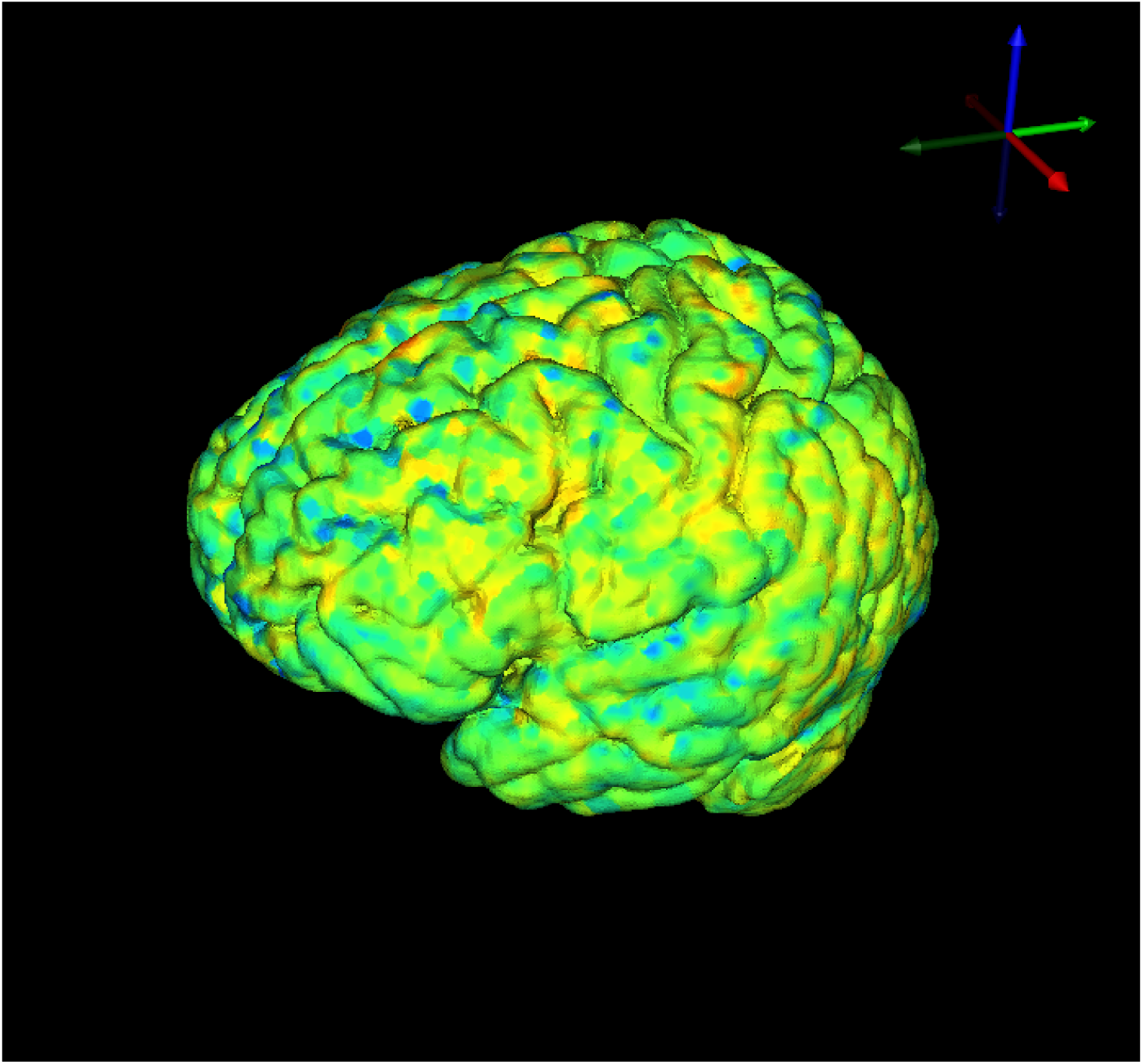
fMRI data mapped onto the cortical surface mesh.

### 3.6 EEG Visualization

When using an EEG dataset, the particular application dictates if further processing, filtering, or cutting of the data is necessary. This EEG dataset, taken with 256 electrodes, contained electrodes that require further processing, specifically around the eyes, possibly due to the blinking or rolling of the subject’s eyes. These electrodes can be corrected or removed with further specific processing, such as trilinear interpolation.

**Figure 32:**
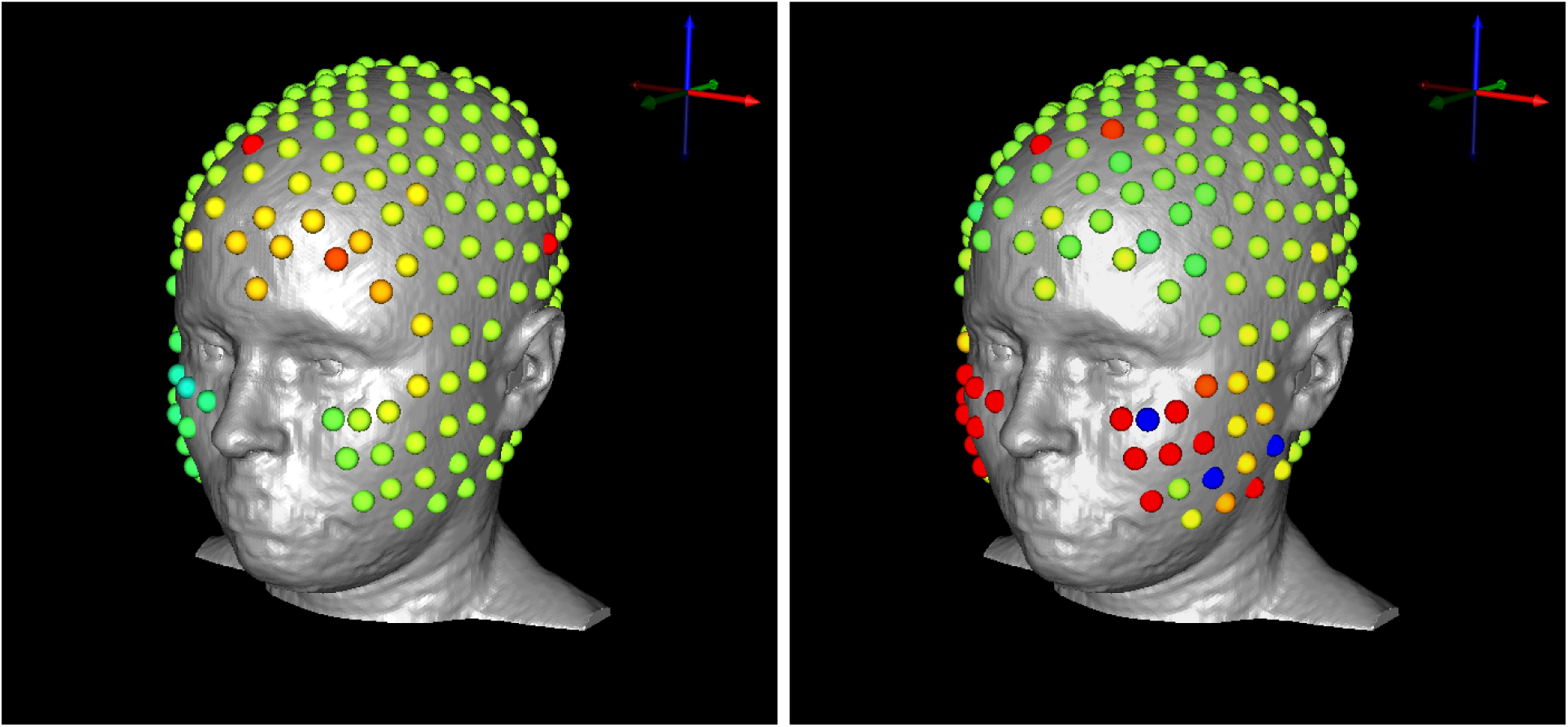
EEG signal visualization with 256 electrodes. Examples of electrodes that require further processing for specific applications shown in red.

The second EEG dataset, taken with 128 electrodes, also contains electrodes that require further processing around the eyes (Figure 33). Since the electrodes do not extend as far down the cheeks as the other dataset, we do not see as many of these electrodes. However, the quality of some time steps was too poor for practical use and can be removed.

**Figure 33:**
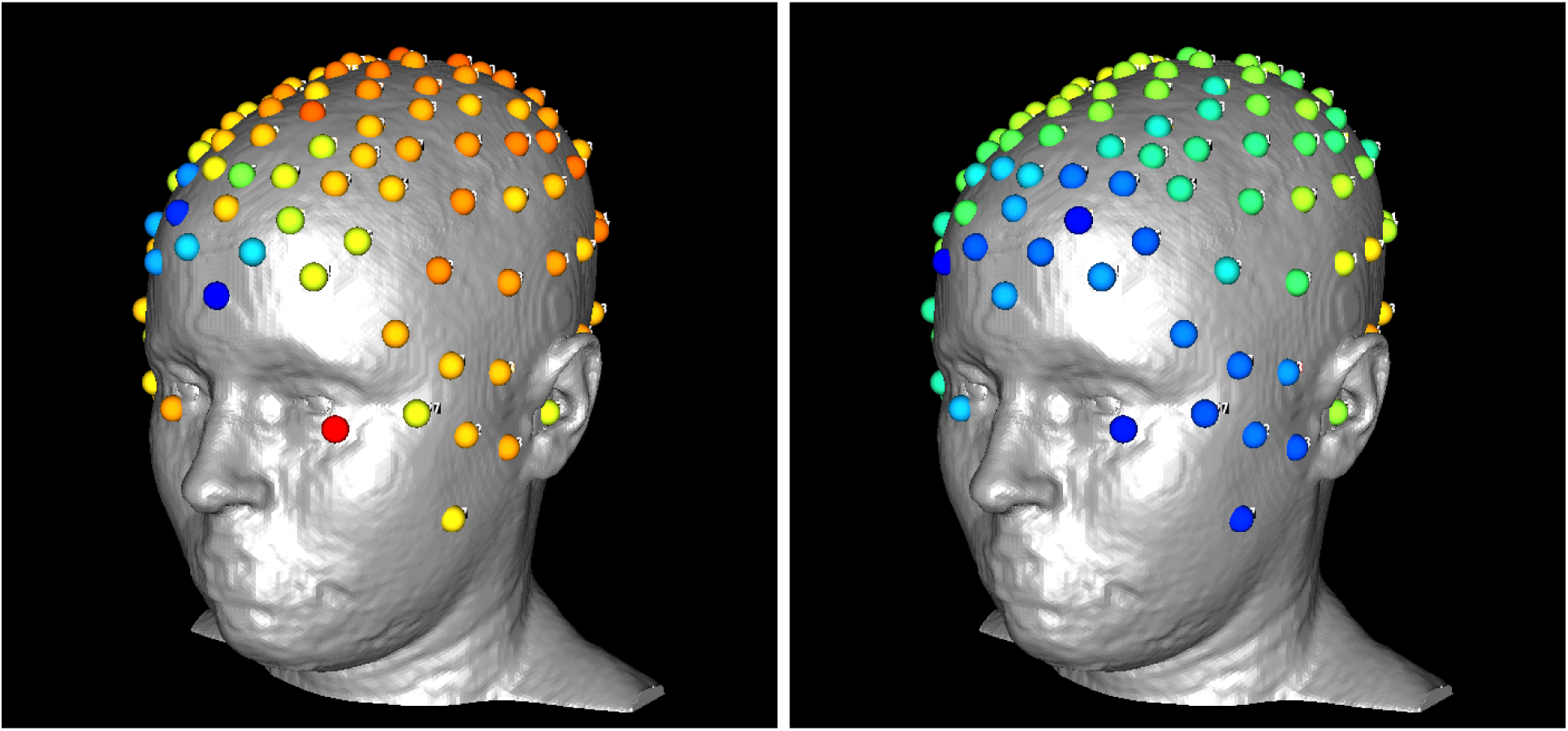
EEG signal visualization with 128 electrodes. Examples of electrodes that require further processing for specific applications shown in red.

The manual registration we applied to the electrodes is also fairly accurate. The spaces for the ears and eyes are where we expected them, and fit the surface of the head mesh without any gapping or shifting.

### 3.7 Future Investigations

Future investigations based on this pipeline include finding more appropriate decimation algorithms for three-dimensional tetrahedral finite element meshes to further reduce the mesh size; more exact sinus and skull segmentation methods to improve the appearance and accuracy of these layers; and more robust registration techniques, which will provide a better transformation matrix for moving images to DTI coordinate space, especially for fMRI data. Additions to this dataset could include more methods to incorporate fMRI data into forward and inverse simulations and more specific processing of EEG data for different applications, resulting in more accurate and realistic visualizations.

## 4 Conclusion

In this project, we have described a comprehensive pipeline to build an inhomogeneous, anisotropic head and brain model based on human data of multiple image modalities for use in electroencephalography with an emphasis on forward and inverse problem research, as well as visualizations of functional MRI and EEG data. Along with the pipeline, we have released the example subject data as open-source to enable other scientists to have a starting point and a straightforward path for further research. The high-resolution, multi-image dataset is available in parts for those who want to use only specific aspects of the project.

The presented female, open-source, and multi-modal dataset will facilitate new research in previously unexplored domains by having a readily available, workable head model that can be used in many simulation applications. Additionally, the pipeline can serve as a template for developing new or specialized image-based head-modeling pipelines.

## 5 Acknowledgements

Imaging and electroencephalography for this project were done at Electrical Geodesics Inc. at the University of Oregon. This project was supported by the National Institute of General Medical Sciences of the National Institutes of Health under grant number P41 GM103545-18.

## Appendix A

The pipeline described in this paper required the use of many software packages and tools. In this appendix, we give more information on how to effectively use these tools, specifically for this project and dataset. We have also included images of registration, tensor field creation, simulation, and visualization networks in SCIRun as a preview to the files in the database.

### A.1 DWI Distortion Correction

#### A.1.1 FSL Total Readout Time

Two parameters are frequently required to calculate and apply field maps to correct distortions: the effective echo spacing and the total readout time for an EPI sequence. We used effective echo spacing, rather than the actual echo spacing, in order to include the effects of parallel imaging, phase oversampling, etc. We defined effective echo spacing as:

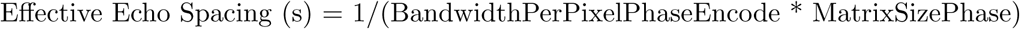

The total readout time (FSL definition) was:

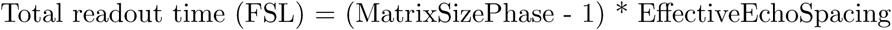

Total readout time, which is a necessary input for using FSL’s topup and eddy tools for DWI image correction, can be obtained by using MRIConvert. Details on using MRIConvert are in Section A.1.2.

#### A.1.2 MRIConvert

The software package MRIConvert provided acquisition information about the dicom series, as well as converted the MRI to NiFTI format, including effective echo spacing and total readout time.^27^

To obtain this information, we loaded the dicom series for either DWI acquisition. We selected “Options” to ensure the DWI was saved in NiFTI format. We then selected “Convert All” to save all the files in the output directory specified upon opening MRIConvert. The text file included the FSL-defined total readout time, which was contained in the acquisition parameter file in seconds as a unit. MRIConvert also output the b-values and b-vectors files, which were the same for both the DWI AP and DWI PA scans. The last input file required was an “index.txt” file, which contained one column with 65 rows (for 64 directions plus the b0 image) of 1’s.

**Figure 34:**
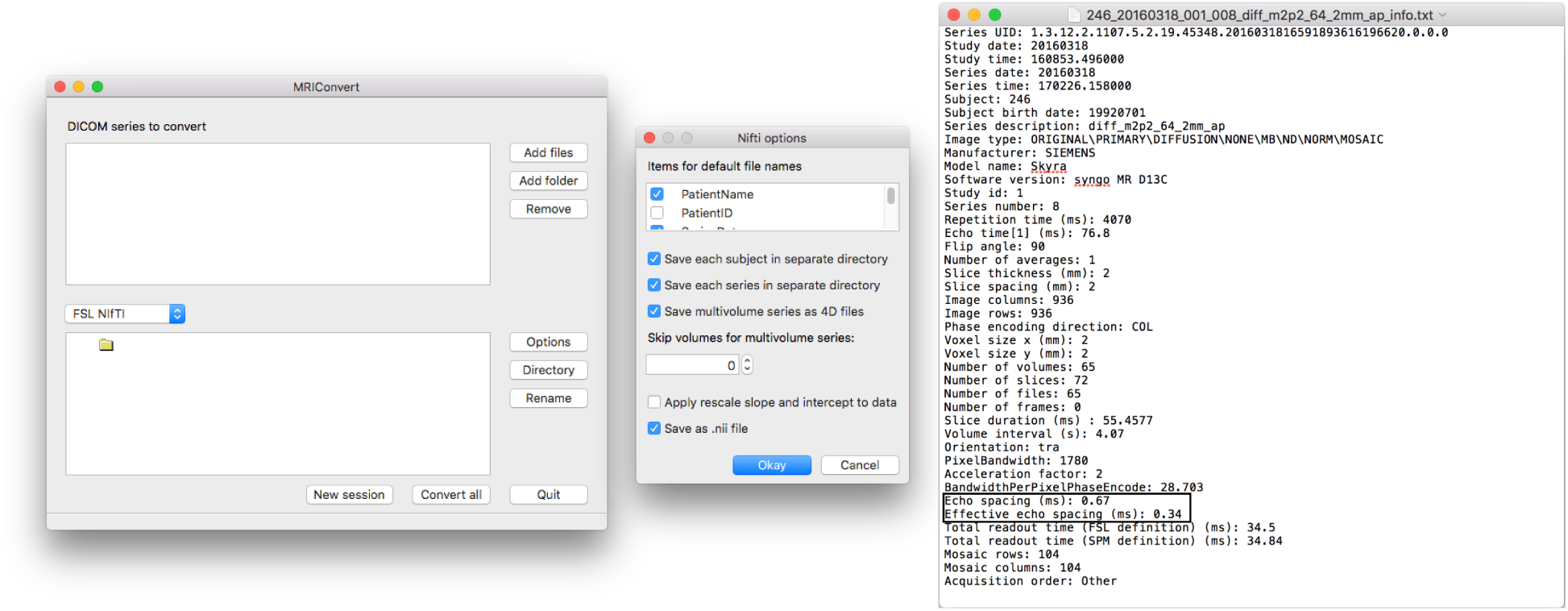
MRIConvert *(left)* with options *(middle)* and output *(right)*.

**Figure 35:**
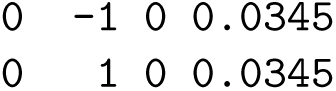
Acquisition parameters text file.

#### A.1.3 FSL’s Topup and Eddy Command Line Tools

We made a separate folder for topup results that included the following files: the acquisition parameters file, the index file, b-values, b-vectors, and the DWI AP and DWI PA files. To run topup, we renamed the DWI AP image DWI up and the DWI PA image DWI down. We renamed the b-values and b-vectors dwi.bval and dwi.bvec, respectively. After all files were in place, we executed the following command line commands:

~~~
fslroi DWI\_up b0\_up 0 1
fslroi DWI\_down b0\_down 0 1

fslmerge -t both\_b0 b0\_up b0\_down

topup --imain = both\_b0 -- datain = acq_params . txt -- config = mine . cnf --out = topup\_results
applytopup --imain =b0\_up, b0\_down -- inindex =1, 2 -- datain = acq_params . txt
           --topup = topup\_results --out =b0\_hifi

bet b0\_hifi b0\_hifi\_brain -m -f 0.2
eddy --imain = DWI\_up --mask =b0\_hifi\_brain\_mask --index = index .txt --acqp = acq_params .txt
     --bvecs =dwi . bvec --bvals =dwi. bval --fwhm =0 --topup = topup\_results --flm = quadratic
     --out = eddy\_unwarped
~~~

By running these commands, we first obtained the b0 image, which is the baseline image used for calculating field maps for both encoding directions. Then the two b0’s were merged together into one file, topup and eddy were applied for distortion correction, and ‘bet’ was applied for brain extraction. The distortion corrected file was named “eddy unwarped.nii.”

### A.2 DTIFIT

To use DTIFIT, we first opened FSL, and then we chose “FDT Diffusion,” followed by “DTIFIT Reconstruct diffusion tensors” in the drop-down menu. We chose the input files manually; Table 4 lists the files selected.

**Table 4:** DTIFIT Input Files

|  |  |
| --- | --- |
| Diffusion weighted data | eddy_unwarmed.nii |
| BET binary brain mask | b0_hifi_brain_mask.nii |
| Output basename | desired output location |
| Gradient directions | dwi.bvec |
| b values | dwi.bval |

DTIFIT output the eigenvalues (named L1, L2, and L3) and the eigenvectors (named V1, V2, and V3) for the diffusion tensor field. We converted the files from NiFTI format to NRRD format using ITK-SNAP.^28^ These are the input files for the SCIRun network in Figure 36.

### A.3 NiFTI Toolbox for fMRI Preprocessing

After running the fMRI data through the fcon pipeline, we converted the preprocessed fMRI data from four-dimensional to two-dimensional. We opened “rest.nii” in Matlab using the “load nii(‘rest.nii’)” function within the NiFTI toolbox.^29^ We then resized the four-dimensional “img” variable into a two-dimensional variable for use in SCIRun using Matlab’s “resize” function.

### A.4 EEG Data Matrix in MATLAB

The EEG data was output in an .edf file format. We calculated the EEG signals matrix using a Matlab script called “edfRead.m.”^30^ To run this script, we called “[hdr, record] = edfread(fname).” The variable ‘record’ contained the EEG signals.

### A.5 Networks

**Figure 36:**
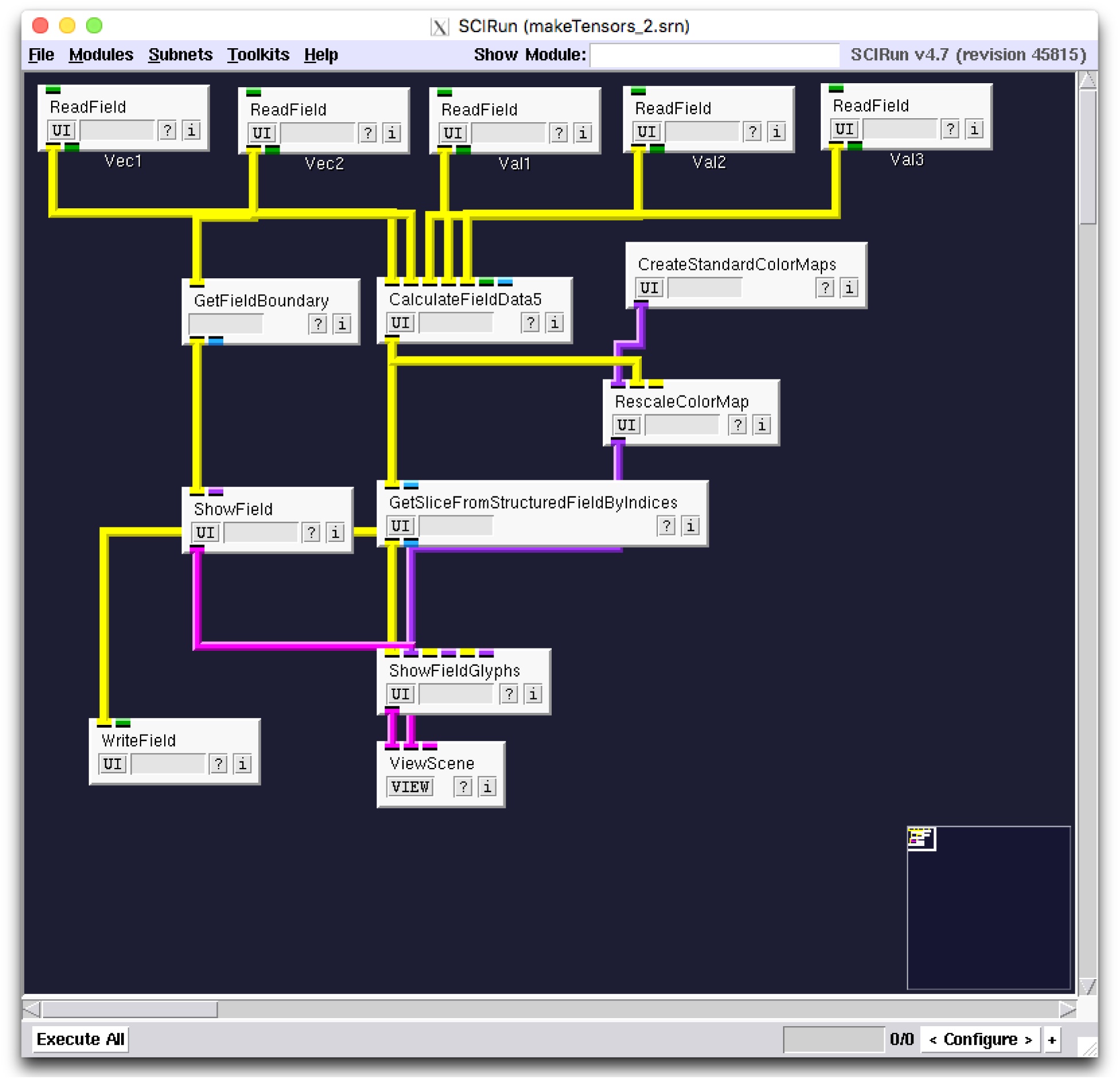
SCIRun network to build and visualize diffusion tensor dataset.

**Figure 37:**
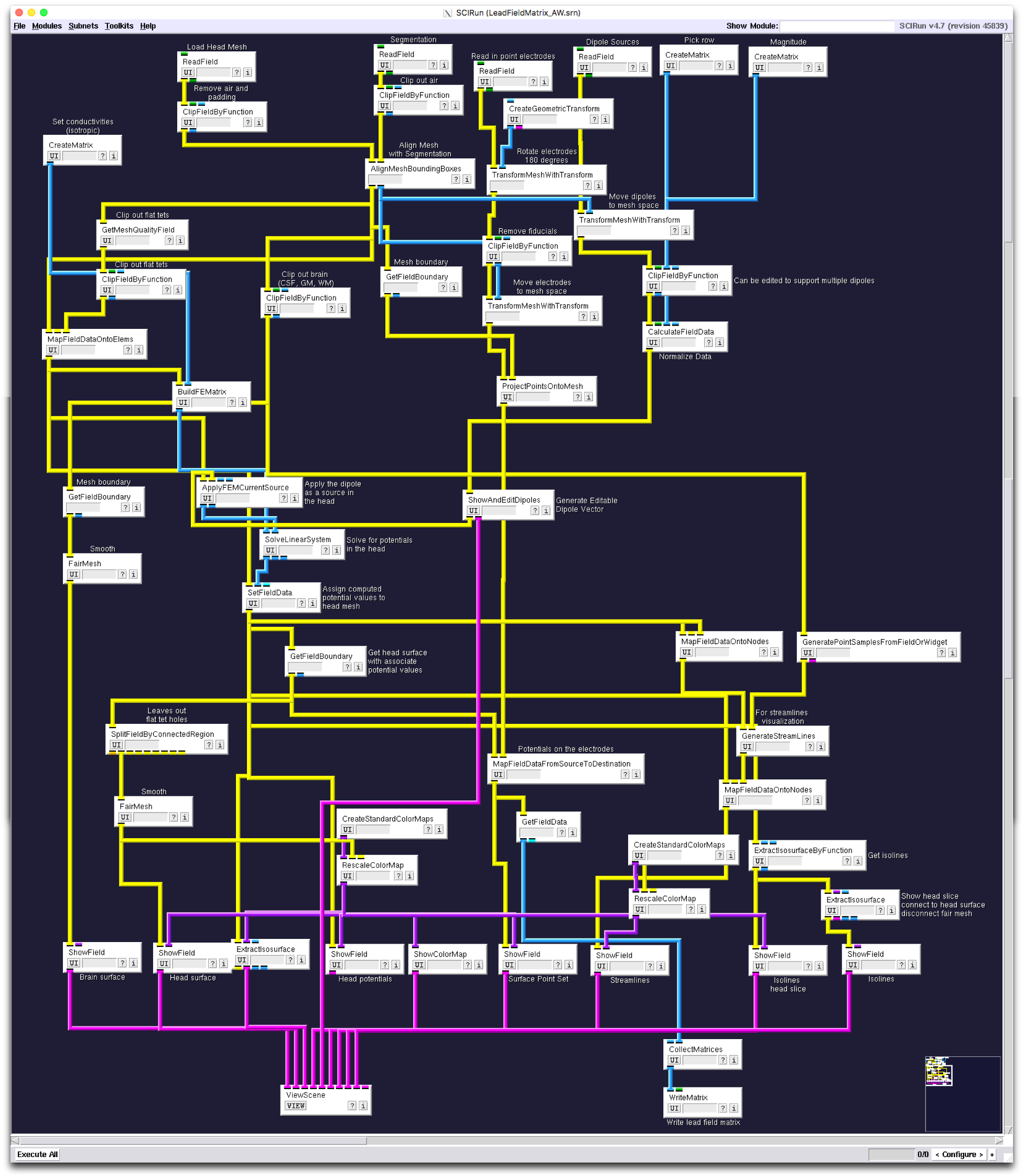
SCIRun network for isotropic forward problem, including registration, visualization, and flat tetrahedra removal.

**Figure 38:**
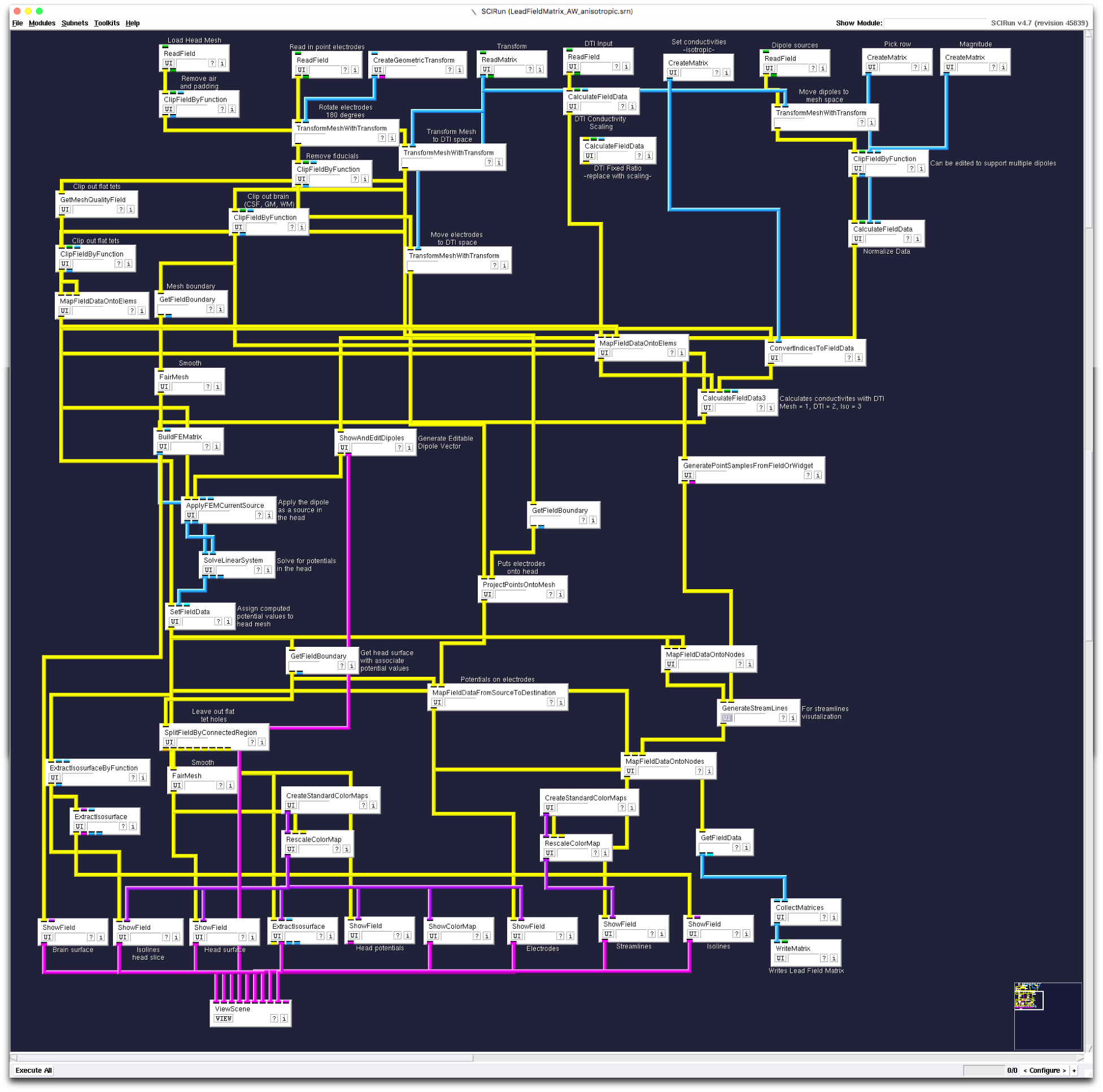
SCIRun network for anisotropic forward problem, including registration, visualization, and flat tetrahedra removal.

**Figure 39:**
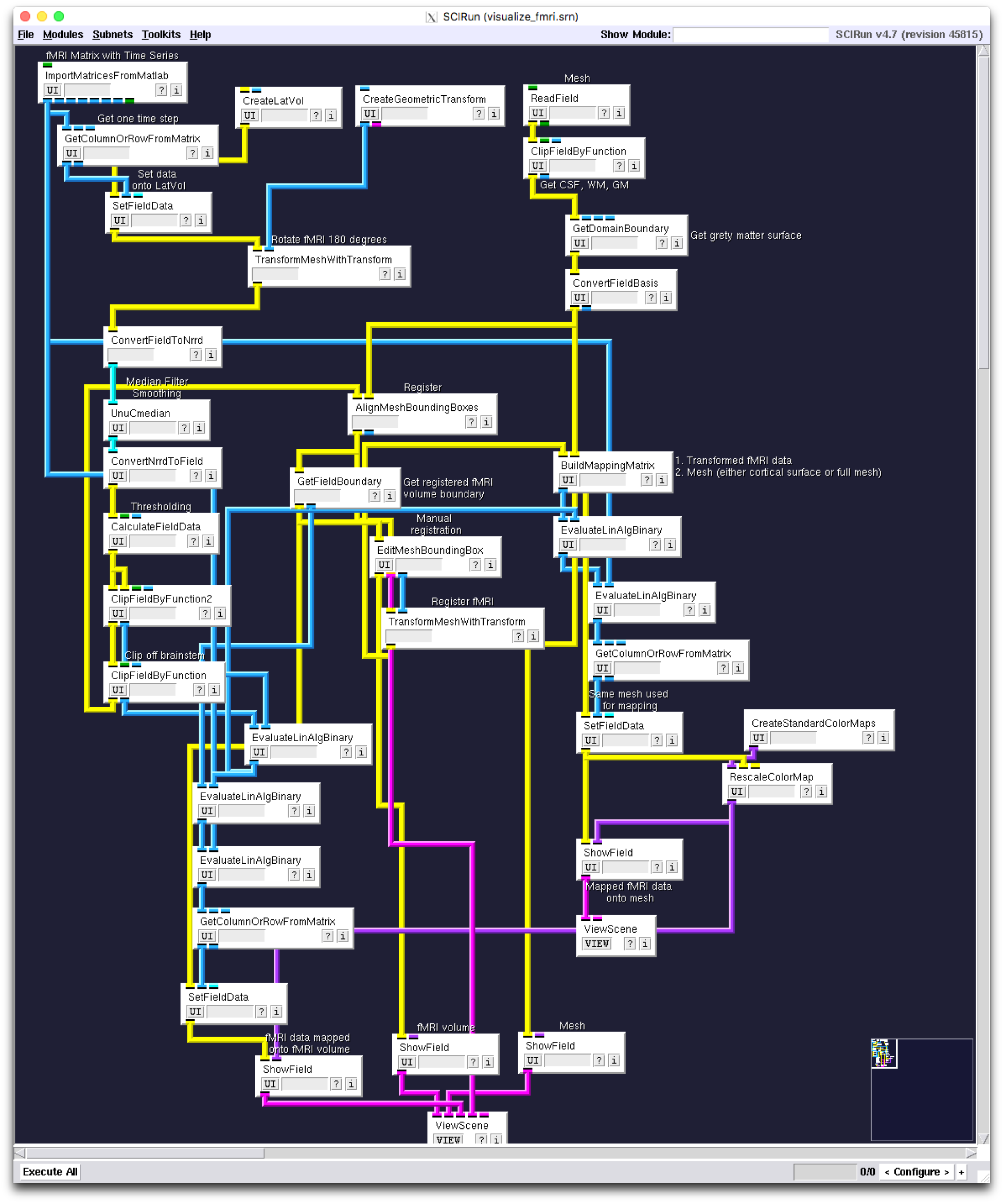
SCIRun network for fMRI registration and visualization.

**Figure 40:**
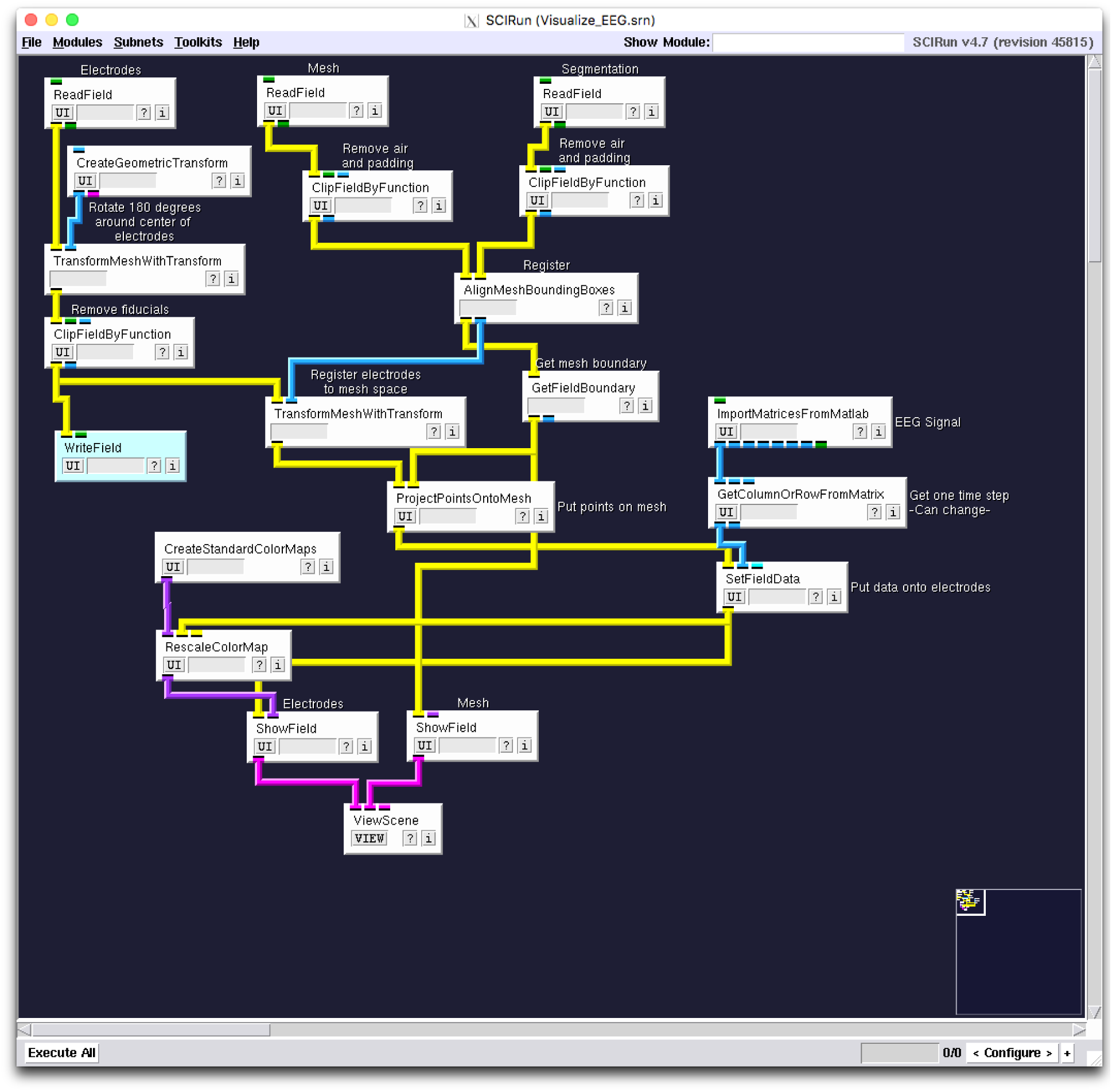
SCIRun network for EEG registration and visualization.

